# Molecular determinants of dynamic protein-protein interactions in the functional cycle of the membrane protein DsbD

**DOI:** 10.1101/2022.09.07.506916

**Authors:** Lukas S. Stelzl, Paraskevi Kritsiligkou, Ahmad Reza Mehdipour, Andrew J. Baldwin, Stuart J. Ferguson, Despoina A. I. Mavridou, Mark S. P. Sansom, Christina Redfield

## Abstract

Molecular recognition is of central importance in biology. The molecular determinants shaping recognition of one protein domain by another are incompletely understood, especially in the context of the complex function of molecular machines. Here, we combine NMR experiments and molecular dynamics simulations to elucidate the determinants of recognition of the C-terminal (cDsbD) domain of the transmembrane reductant conductor DsbD by its cognate partner, the N-terminal domain of the protein (nDsbD). As part of the natural cycle of this oxidoreductase, which effectively transfers electrons from the cytoplasm to the periplasm of Gram-negative bacteria, cDsbD and nDsbD toggle between oxidised and reduced states, something that modulates the affinity of the domains for each other and prevents otherwise unproductive reactions. We find that the redox state of cDsbD determines the dissociation rate of cDsbD-nDsbD complexes. Molecular dynamics simulations demonstrate how the redox-state of the active site determines the stability of inter-domain hydrogen bonds and thus the dissociation rate. AlphaFold modelling and atomistic molecular dynamics simulations of full-length DsbD in a realistic bacterial membrane again highlights the close proximity of the periplasmic domains and the importance of tuning the strength of the interactions of the periplasmic domains to enable electron transfer to cognate periplasmic partners such as CcmG. Our AlphaFold models are consistent with *in vivo* functional assays of DsbD mutants, which together help to reveal for the first-time a putative binding site for thioredoxin on the cytoplasmic side of DsbD.

## Introduction

The association and dissociation of biomolecules is central to biology,^1^ but the molecular determinants of protein-protein association remain to be fully elucidated. Advances in artificial intelligence (AI) such as AlphaFold^2,3^ complement experiments in revealing the different complexes a protein might form as part of its function. Nonetheless how protein-protein association is mediated by subtle biochemical cues^4^ in the functional cycles of proteins awaits elucidation. NMR spectroscopy and, in particular, the relaxation dispersion method^5,6^ has revealed intermediates in the association reactions of proteins.^7–9^ Molecular dynamics (MD) simulations also can, in favourable cases resolve protein-protein association and dissociation with atomic resolution.^10–13^ Simulations are particularly powerful in tandem with experiments,^14^ highlighting aspects of molecular recognition not directly accessible otherwise.^1,15^

Even subtle chemical differences can play decisive roles in recognition at the molecular scale as has been shown for the interaction of the periplasmic N- and C-terminal domains of the transmembrane protein DsbD (nDsbD and cDsbD, respectively)^16–18^(Fig. 1A). DsbD functions as a reductant conductor that transfers reducing equivalents from the cytoplasm to the periplasm of Gram-negative bacteria, an otherwise, oxidising environment. Electrons are required for the refolding of proteins with disulfide bonds, the protection against oxidative stress and cytochrome *c* maturation, a crucial protein for bacterial respiration. During the catalytic cycle of this protein, two electrons and two protons are transferred across the three domains of DsbD, spanning the inner membrane and extending into the periplasm, in a series of thiol-disulfide exchange reactions (Fig. 1B). During this thiol-disulfide cascade protein domains must form specific complexes, permitting electron transfer via a covalent, disulfide-bonded intermediates (Fig. 1C). After electron transfer from cDsbD to nDsbD, as part of its natural function, the thermodynamic stability of the complex decreases. By destabilizing this complex underpinning electron transfer the redox-state dependence of the interaction does not only prevent the futile but thermodynamically feasible transfer of electrons from nDsbD back to cDsbD, but also frees up the two domains for further reductant transfer steps. cDsbD can then receive reductant from the trans-membrane domain of DsbD (tmDsbD), whereas nDsbD can then distribute electrons to different pathways, acting as a periplasmic “redox hub”.^19^ The redox-state dependence of the interaction affinities may enhance turnover during DsbD function. Like with other thiol-disulfide oxidoreductases, electron transfer in DsbD is mediated by mediated by thiol-disulfide exchange reactions, involving conserved pairs of cysteine residues; nDsbD and cDsbD cycle between redox isoforms with a pair of cysteine thiols (reduced) or a disulfide bond (oxidised) in their active site. In nDsbD_ox_, the presence of the disulfide bond in its active site leads to a dynamic structural equilibrium which includes the transient cap loop opening required for formation of a complex with cDsbD; this loop opening is not observed in nDsbD_red_ in the absence of its cognate binding partners.^18^

**Figure 1:**
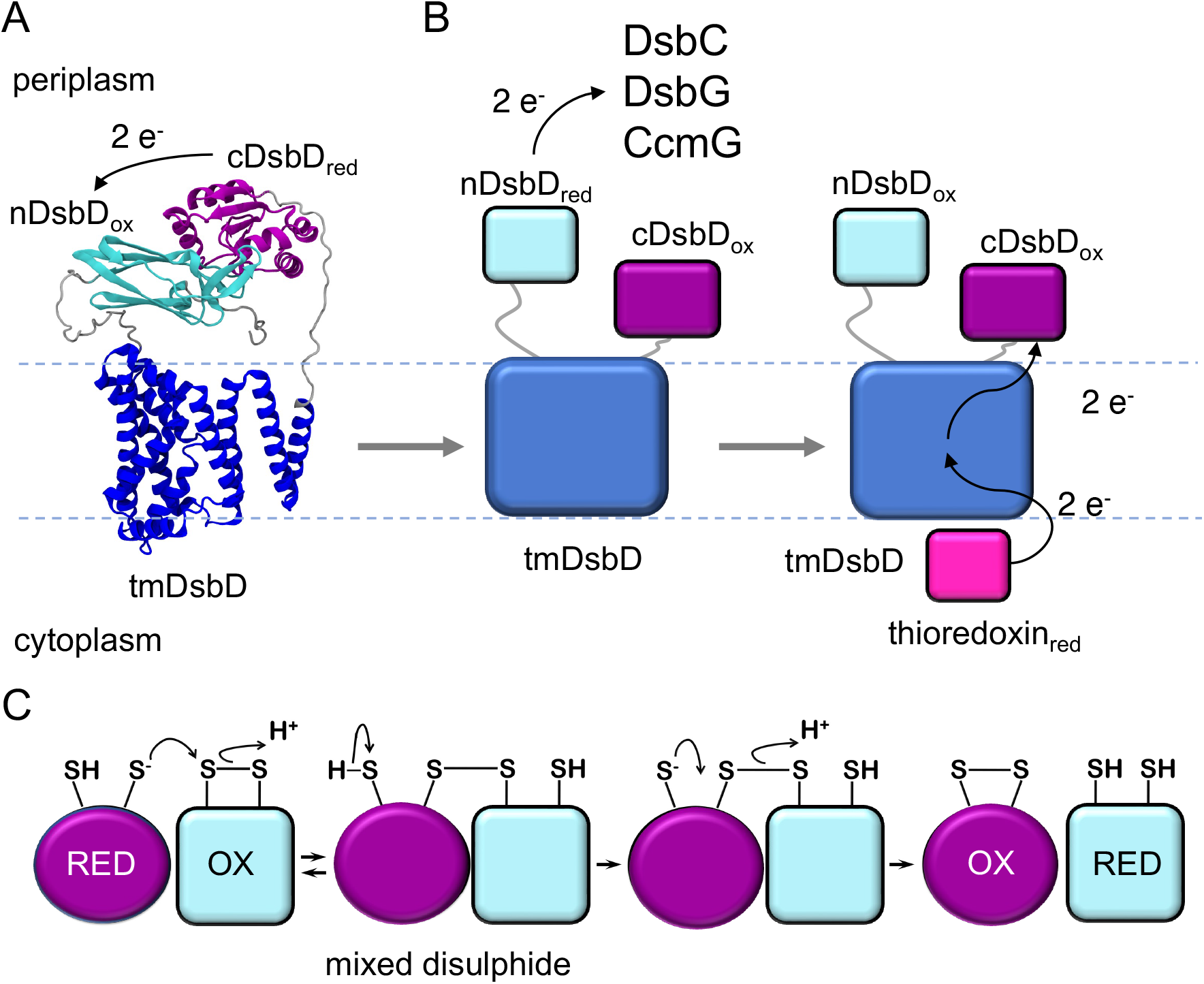
The C-terminal domain of DsbD, cDsbD, receives reductant (2e^−^ and 2 H^+^) from the transmembrane domain of DsbD. cDsbD passes on reductant to nDsbD. A) Full-length model of DsbD generated with AlphaFold. Possible location of the membrane is indicated. Electron transfer from cDsbD_red_ to nDsbD_ox_ is indicated. B) nDsbD_red_ supplies electrons to DsbC, DsbG, and CcmG. cDsbD_ox_ receives electrons from thioredoxin via the transmembrane domain of DsbD (tmDsbD). C) DsbD transfers electrons via thiol-disulfide exchange reactions involving an inter-molecular disulfide bond which necessitates the formation of sterospecific complexes of the domains involved

The AlphaFold model^2,20^ of DsbD (AF-P36655-F1) provides a first atomically-detailed static view of the 3D structure of full-length DsbD. The model shows cDsbD and nDsbD bound to each other (Fig. 1A), as observed in the previous X-ray structure of a covalent mixed-disulfide complex, ^21^ but it does not explain how the domains associate or dissociate so that cDsbD can receive electrons from thioredoxin via the transmembrane domain of DsbD. The two domains also need to dissociate so that nDsbD can supply electrons to partners such CcmG of the cytochrome *c* maturation pathway. Linkers of about 20 amino acids connect nDsbD and cDsbD to the transmembrane domain. While the linkers are long enough for the domains to fully detach from the transmembrane domain, it is clear that high-local concentrations as well as the proximity of the membrane will shape the interactions of the periplasmic domains. Importantly, the transmembrane strucure of the related protein CcdA has been resolved by NMR,^22^ and this can be used to further inform studies of full-length DsbD and its function.

In particular, the AlphaFold model of DsbD does not resolve how the redox-state of the cysteines in the active site of cDsbD determines the strength of cDsbD-nDsbD interactions nor does it provide insight into the kinetics of complex formation and dissociation. AlphaFold has not been specifically tested and validated for predicting the effects of small changes such as a point mutations or changes in the redox-state of cysteines on protein-protein interactions. Redox-state and resulting change in the *pK*_*a*_*s* of cDsbD residues may affect the surface of electrostatics of cDsbD ^16,23^ which in turn, could affect the equilibrium of its reaction with nDsbD, for example, by changing the association rate *k*_on_. Alternatively, the redox state of cDsbD affects its hydrogen bonding network^24^ which could then influence the stability of the complex, i.e., the *k*_*off*_. The stability might also be influenced more directly by the redox state of the two interacting domains. Two thiol groups fill more space than a disulfide bond. Packing constraints introduced by reducing a disulfide bond to two thiols may determine whether the hydrogen bonds between cDsbD and nDsbD can be maintained as emphasised by energy minimisation *in vacuo*.^23^ Such a direct role of the active site cysteines resembles the situation found for nDsbD.^18^ Many proteins engaging in electron transfer reactions dissociate quickly with *k*_*off*_ in the range of 1000 s^*−*1^. Besides understanding how the oxidoreducatese prevents back reactions and increases turnover, kinetic measurements will give insight into the activity of DsbD. The rate for reductant transfer between cDsbD and nDsbD was found to be off the order of 1 × 10^6^ M^*−*1^ s^*−*1^.^21^ It will be interesting to see whether reductant transfer between the periplasmic domains of DsbD is limited by the association rate and would thus be “catalytically perfect”.^1^

To understand the molecular determinants of molecular recognition, we study how a subtle chemical modification, as part of its natural function, shapes the kinetics of complex formation between cDsbD and nDsbD. With NMR relaxation dispersion experiments which permit an in-depth characterisation of protein-protein interactions which would be difficult to measure otherwise, we find that the redox state of cDsbD modulates its dissociation rate. Our molecular dynamics simulations highlight how the redox state of the active site affects the stability of its complexes. To put these findings into the context of the functional cycle of DsbD, we use AlphaFold to model different functional intermediates of DsbD and its cognate binding partners. These structural models are consistent with *in vivo* experiments on the function of DsbD mutants, which together also suggest a binding mode of cytoplasmic thioredoxin to the transmembrane domain of DsbD to resupply DsbD with electrons.

## Results

### NMR relaxation dispersion experiments on the interaction of C461A-cDsbD with oxidised nDsbD

The association between C461A-cDsbD and oxidised nDsbD was followed by NMR relaxation dispersion experiments. Non-reactive C461A-cDsbD mimics the reduced form of the protein and therefore its complex with nDsbD resembles the productive complex prior to electron transfer. Measurement of relaxation dispersion effects was done using ^15^N labelled C461-cDsbD because, in the absence of nDsbD_ox_, C461A-cDsbD shows no dispersion effects (Fig. 2). By contrast, nDsbD_ox_ does show dispersion effects in the absence of a binding partner and so is not suitable for monitoring the effects of interaction with cDsbD.^18^

**Figure 2:**
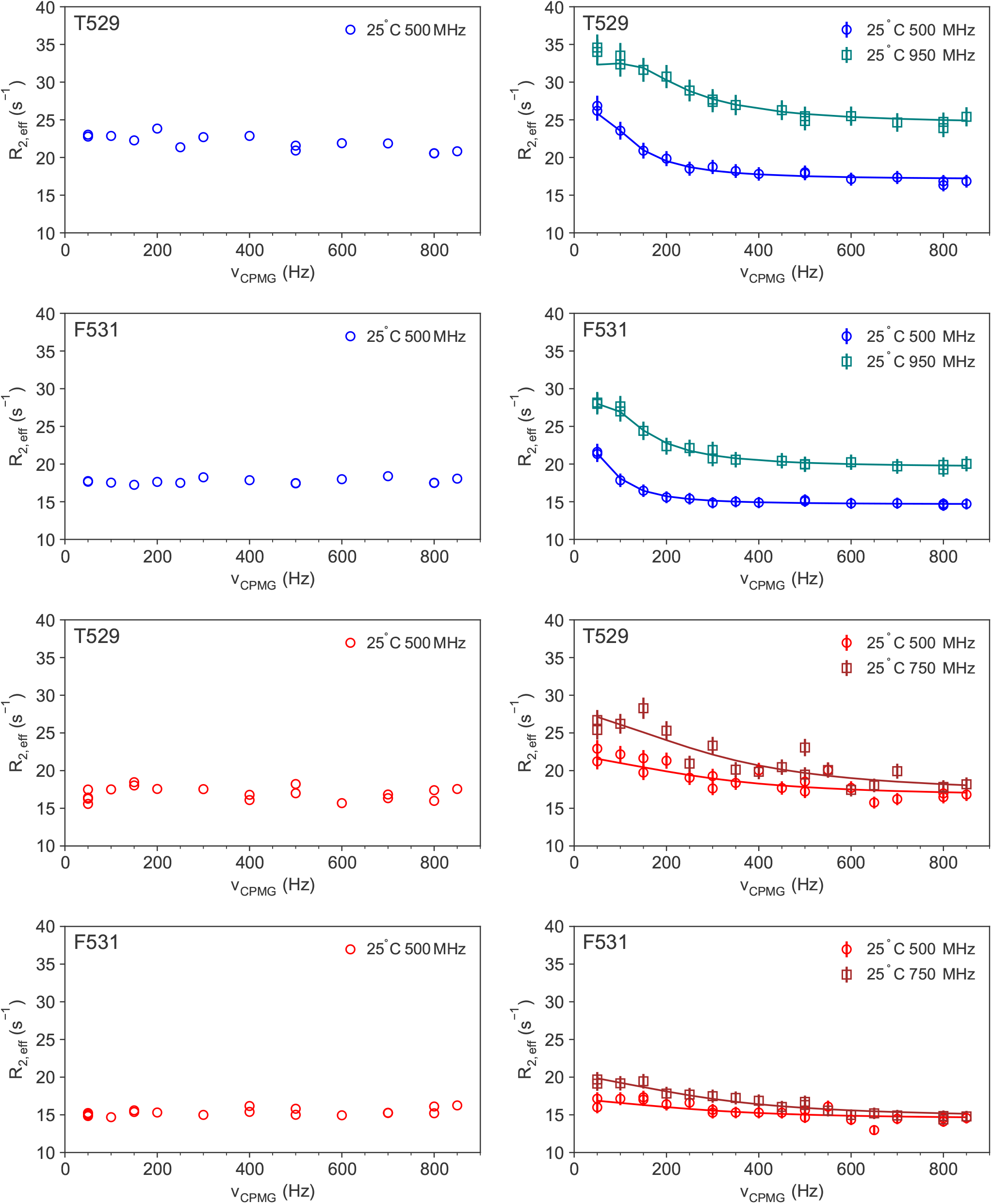
NMR relaxation dispersion curves for residues T529 and F531 in the absence and presence of nDsbD. The top two rows shows data for ^15^N C461A-cDsbD in absence of nDsbD_ox_ (left) and in presence of substoichmetric amounts of nDsbD_ox_ (right). The bottom two rows shows data for ^15^N cDsbD_ox_ in absence of nDsbD_ox_ (left) and in presence of substoichmetric amounts of nDsbD_ox_ (right).

A small amount of unlabelled nDsbD was added to ^15^N labelled C461-cDsbD. 1D NMR experiments showed that there was about 25-fold more cDsbD than nDsbD in the sample (Fig. S1). This should provide a good trade-off between detecting all key residues, which experience significant line broadening at higher concentrations of nDsbD, and having a large enough population of the minor, nDsbD-bound, state to observe dispersion. Good relaxation dispersion curves were indeed obtained under these conditions as shown Fig. 2 and Fig. S4. Inspection of the relaxation dispersion curves demonstrated that the reaction is in relatively slow exchange, as expected from a previous NMR study.^23^ Representative relaxation dispersion curves are shown for T529 and F531. These curves level off at refocusing fields around 300 Hz. As the curves appeared to have levelled-off completely, good fits should be obtained. The relaxation dispersion curves for the association of C461A-cDsbD with nDsbD were analysed using CATIA^25–27^ A single global exchange process was found to account for the experimental curves of the 22 residues at two magnetic fields (Table 1). Residues that showed significant dispersion |Δppm| *>* 1 ppm are in the interaction interface with nDsbD as shown in Fig. 3A. The exchange is slow as expected, with a rate *k*_*ex*_ of 220 s^*−*1^. About 4 % of cDsbD is in complex with nDsbD. The population of the bond state is consistent with independent estimates. nDsbD-bound cDsbD is visible in 1D ^1^H spectra. Fitting a Lorentzian to the peaks for the bound and free protein suggested that roughly 2 % of cDsbD would be in the complex. The population of bound cDsbD from the fits also compared favourably to the estimate based on the dissociation constant (*K*_*d*_ = 86 μM) and the concentrations of cDsbD and nDsbD, which suggested that about 3 % of cDsbD would be bound to nDsbD, clearly in the same range as the bound fraction determined from relaxation dispersion fits.

**Table 1:** **Relaxation dispersion fit parameters for the association of cDsbD with oxidised nDsbD and the macroscopic on and off-rates determined from them. The fit parameters for the association reactions with C461A-cDsbD, which mimics the reduced isoform, and with oxidised cDsbD are given in the table**

| | C461A-cDsbD | oxidised cDsbD | oxidised cDsbD ($p_B$ fixed) |
| --- | --- | --- | --- |
| $k_{ex}$ | $220 \pm 26\text{ s}^{-1}$ | $2500 \pm 200\text{ s}^{-1}$ | $1720 \pm 130\text{ s}^{-1}$ |
| $p_B$ | $4 \pm 0.4\%$ | $2.9 \pm 0.3\%$ | $3.0\%$ |
| $k_{off}$ | $210 \pm 33\text{ s}^{-1}$ | $2430 \pm 200\text{ s}^{-1}$ | $1670 \pm 120\text{ s}^{-1}$ |
| $k_{app}$ | $10 \pm 1\text{ s}^{-1}$ | $70 \pm 10\text{ s}^{-1}$ | $50 \pm 5\text{ s}^{-1}$ |
| $k_{on} = \frac{k_{off}}{K_d}$ | $2.4 \times 10^6\text{ M}^{-1}\text{ s}^{-1}$ | $3.0 \times 10^6\text{ M}^{-1}\text{ s}^{-1}$ | $2.1 \times 10^6\text{ M}^{-1}\text{ s}^{-1}$ |

**Figure 3:**
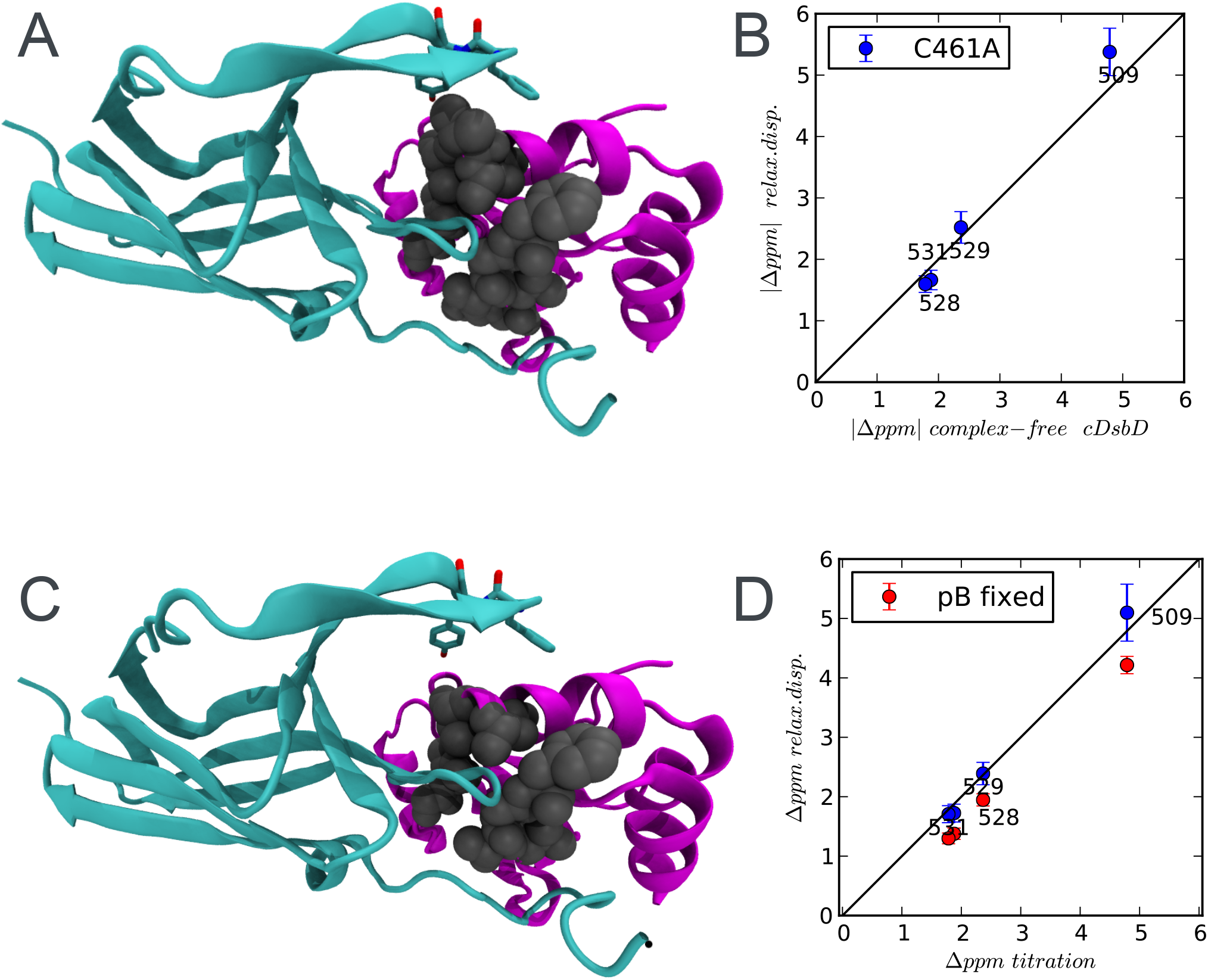
Chemical shift differences for complex formation determined by NMR relaxation dispersion experiments. (A) The residues in C461A-cDsbD that show significant dispersion in the association reaction with nDsbD are highlighted. Residues in gray, G509, L510, R527, V528, T529, G530, F531, C461, V462, A463 and C464, show |Δppm| *>* 1.0 ppm. CDsbD (1VRS E) is shown in magenta and nDsbD (1VRS E) in cyan. (B) Correlation between absolute chemical shift differences from relaxation dispersion C461A-cDsbD-nDsbD_ox_ and covalent cDsbD-nDsbD complex. For G509, V528, T529, and F531 the bound shifts could be determined from the covalent complex with confidence. (C) cDsbD_ox_ residues, C461, A463, C464, G509, L510, V528, T529, G530 and F531, for which |Δppm| *>* 1.0 ppm are shown in grey. cDsbD (1VRS E) is shown in magenta and nDsbD (1VRS E) in cyan. (D) Correlation between absolute chemical shift differences from relaxation dispersion of cDsbD_ox_-nDsbD_ox_ and covalent cDsbD-nDsbD complex for G509, V528, T529, and F531.

To validate further the interpretation of the relaxation dispersion experiment, the chemical shift differences between the bound and free forms of cDsbD from CATIA fitting and from previous titration experiments ^23^ were compared. Mavridou *et al*. determined the shift differences for complex formation based on the covalent complex of the two domains. Since the complexes in the two studies are not identical, a good, but not a perfect, correlation would be expected. A good correlation between the chemical shift differences from the two methods was indeed obtained as shown in Fig. 3B. This further confirms the validity of the fit parameters and one can conclude that a single global, two-state, exchange process describes the data well.

Having validated the analysis of the fits, the macroscopic on and off rates for the complex formation between C461A-cDsbD and oxidised nDsbD can be determined (Table. 1) ^8^. The off rate, *k*_*off*_, is given simply by *k*_*off*_ = *p*_*A*_*k*_*ex*_ and is 210 s^*−*1^. Using the dissociation constant previously determined by surface plasmon resonance, *K*_*d*_ =86 μM, an association rate of 2.4 × 10^6^ M^− 1^s^− 1^(Table. 1) was calculated.

### NMR relaxation dispersion experiments of the interaction of oxidised cDsbD with oxidised nDsbD

Relaxation dispersion experiments were also recorded for the association of oxidised cDsbD with oxidised nDsbD. By changing the oxidation state of cDsbD but not of nDsbD, we can directly probe the influence of the oxidation state of cDsbD on the formation and dissociation of its complex with nDsbD. Previous experiments have highlighted that adding sub stochiometric amounts of oxidised nDsbD to oxidised cDsbD leads to significant peak broadening and the peak of V462 becoming unobservable, almost as soon as ligand was added. Upon adding unlabelled nDsbD to ^15^N labelled cDsbD_ox_ at a ratio of 1:17, as determined by 1D NMR measurements (Fig. S2), we find that that key residues such as G509 that broaden significantly at higher nDsbD concentrations ^23^ were still observable in 2D HSQC spectra.

For multiple residues of cDsbD clear relaxation dispersion effects were observed (Fig. 2 and Fig.S5). Like C461A-cDsbD, cDsbD_ox_ does not show dispersion effects in the absence of nDsbD (Fig. 2). Here the curves for T529, which is located in the cDsbD active site as well as in the nDsbD binding interface, and F531 which is also in the binding site are shown (Fig. 3). For V462, which a key residue stabilising complexes of cDsbD and nDsbD, noisy dispersion data was obtained at magnetic-field strength of 500 MHz. At 750 MHz peaks could not be assigned with confidence for this residue, likely due to extensive exchange broadening. Overall the dispersion curves for ^15^N cDsbD_ox_ interacting with nDsbD_ox_ suggested a much faster exchange process than for the association of C461A-cDsbD with nDsbD_ox_. The curves only appeared to level-off around 800 Hz. CPMG relaxation dispersion curves for such fast exchange processes can be challenging to interpret. Furthermore, the exchange process was likely not fully refocused at the *ν*_*CPMG*_ frequencies up to 850 Hz, that could routinely be accessed on our spectrometers. The incomplete refocusing will somewhat complicate the data analysis and will increase errors in the fit parameters.

We repeated the relaxation dispersion experiments of the interaction of ^15^N cDsbD_ox_ with nDsbD_ox_ at lower temperature of 15 °C, attempting to slow down the exchange between free and bound ^15^N cDsbD_ox_. Interestingly, the shape of the relaxation dispersion curves does not indicate any slowing of the exchange reaction (Fig. S3). The exchange from the major population of free oxidised cDsbD to the minor state of cDsbD_ox_-nDsbD_ox_ likely does not involve an enthalphic barrier.

All residues for which clear dispersion effects were identified were included in the fitting using CATIA. A global exchange process was assumed. The fitting of data recorded at two magnetic field strength for 18 residues (Fig. S5) suggested a process with an exchange rate of 2500 s^*−*1^ and a population of the bound state of 2.9% (Table 1). Based on the concentrations of the two domains and the previously determined dissociation constant, about 3% of cDsbD molecules are expected to be in the complex. As for C461A-cDsbD, residues for which the fit revealed large chemical shift changes between bound and free conformations are in the interface with nDsbD (Fig. 3C). The chemical shift differences between the bound and the free states of cDsbD agree well with the previously determined chemical shift differences for the formation of the complex (Fig. 3D). Considering that both the population of the excited state and the chemical shift differences agree with our previous study it appears that the relaxation dispersion experiments capture the exchange process well. In general *p*_*B*_ and Δppm cannot be determined with confidence from relaxation dispersion curves for such a fast process^28,29^. Nevertheless enough residues have large chemical shift differences between the bound and the unbound state and are outside the fast-exchange limit to enable the determination of *p*_*B*_ and Δppm in the global fitting. Yet, the fitted value for *k*_*ex*_ is rather large, given that curves appeared to level-off around 800 Hz.

To explore whether the exchange is really that fast, the fits were repeated with a more constrained parameter space. For this *p*_*B*_ was fixed to the previously determined value of 3 %. The exchange rate is now 1720 s^*−*1^, slower than *k*_*ex*_ from the unconstrained fits. The chemical shift differences were still reproduced as shown in Fig. 3D, which suggests that overall meaningful parameters can be extracted for this relatively fast exchange process.

The macroscopic on and off rates are also obtained for the association of oxidised cDsbD with oxidised nDsbD. Dissociation rates of 2430 s^*−*1^ and 1670 s^*−*1^ were obtained with *p*_*B*_ either free to vary in the fitting or fixed to 3 %. While there is a considerable difference between the values, it is clear that the dissociation rate is fast, even compared to electron-transferring proteins (*k*_*off*_ = 1000 s^*−*1^) that are known for their fast kinetics ^9^. For both values of *k*_*ex*_ an association rate on the order of 1 × 10^6^ M^*−*1^ s^*−*1^ was determined as shown in Table 1. The association rates for C461A and oxidised cDsbD are of similar magnitude, with a larger value for C461A-cDsbD.

### Computational modelling of the association of cDsbD and nDsbD

To better understand the association of cDsbD_red_ and nDsbD_ox_, we employed atomistic molecular dynamics (MD) simulations, electrostatic calculations and modelling of the association rate, which suggests that the association is not strongly enhanced by electrostatic interactions. By running simulations with no added salt and with 100 mM salt, we investigated whether electrostatic interactions drive the initial encounter of the two domains. Absence of salt should favour protein-protein association driven by electrostatics,^13^ but we did not find evidence for this in our simulations (Supporting information and (Fig S6A,E). In the simulations we observed transient encounter complexes (Fig S6B,F,G,H,I), including long-lived encounter complexes involving the cognate cDsbD-nDsbD binding interface (Fig S6B), but the cap loop remained closed preventing the formation of hydrogen bonds between C109 and L510 and G107 and F531, which stabilise the native complex. Electrostatics calculations show that there is a large negatively charged patch around the cap loop of nDsbD, but there is not a clearly defined complementary positively charged surface on cDsbD, which would accelerate the association by electrostatically steering the two domains towards complex formation (Fig. S6J) Importantly, the full-length AlphaFold model of DsbD implies that the cDsbD-nDsbD complex is formed close to the surface of the membrane. Charged lipids in the membrane would also argue against a functionally role of electrostatic modulation of *k*_on_. In addition, Transition Complex Theory calculations ^30,31^ also suggest that the shapes of cDsbD and nDsbD and their interaction interface determine their association rate and that electrostatics overall does not enhance the association of the two domains (Supporting Information).

### Active-site conformations determine the stability of interactions of the cDsbD-nDsbD complex in MD simulations

MD simulations demonstrate that the active-site structures determine the relative stability of cDsbD-nDsbD complexes. We have simulated the most and least-stable complexes which occur in the physiological electron transfer reaction, to ascertain how differences in the active site affect the interaction interface of cDsbD-nDsbD. The nDsbD_ox_-cDsbD_red_ complex cannot be studied directly with NMR as electron transfer occurs rapidly on time scale of the NMR experiments resulting in mixtures of nDsbD_red_ and cDsbD_ox_. However, with simulations we can study this complex and its inter-domain interactions directly and compare it to other complexes.

The subtle redox-state dependent differences in active-site conformations destablise the inter-domain hydrogen bond between G107 of nDsbD and F531 of cDsbD when nDsbD is reduced. The inter-domain hydrogen bond between G107 and F531 is one of the key interactions stabilizing the complex.^23,32^ The hydrogen bond fluctuates but remains mostly intact, for the relatively stable nDsbD_ox_-cDsbD_red_ complex, which forms before the electron-transfer reaction (Fig. 4A). For the complex of the products of the physiological electron transfer reaction nDsbD_red_-cDsbD_ox_, the hydrogen bond shows considerable fluctuations, breaks and reforms before breaking for a sustained period of time towards the end of the trajectory shown in Fig. 4B. The breaking of the hydrogen bond coincides with the changes in the G107 *ψ* angle to values close to 0 ^°^, flipping the carbonyl of G107 away from the amide group of F531 (Fig. 4B,D). Transient breaks of the hydrogen bond in the simulation of the stable nDsbD_ox_-cDsbD_red_ complex also coincide with excursions of the G107 *ψ* angle to values close to 0 ^*°*^ for instance at 50 ns (Fig. 4A,C). The other two key hydrogen bonds C109 HN to L510 O and C109 O to L510 HN are maintained through the simulations for both the stable and the less stable complexes (Fig. S9). The bidentate interaction likely mutually stabilises these two hydrogen bonds (Fig. 4E,F).

**Figure 4:**
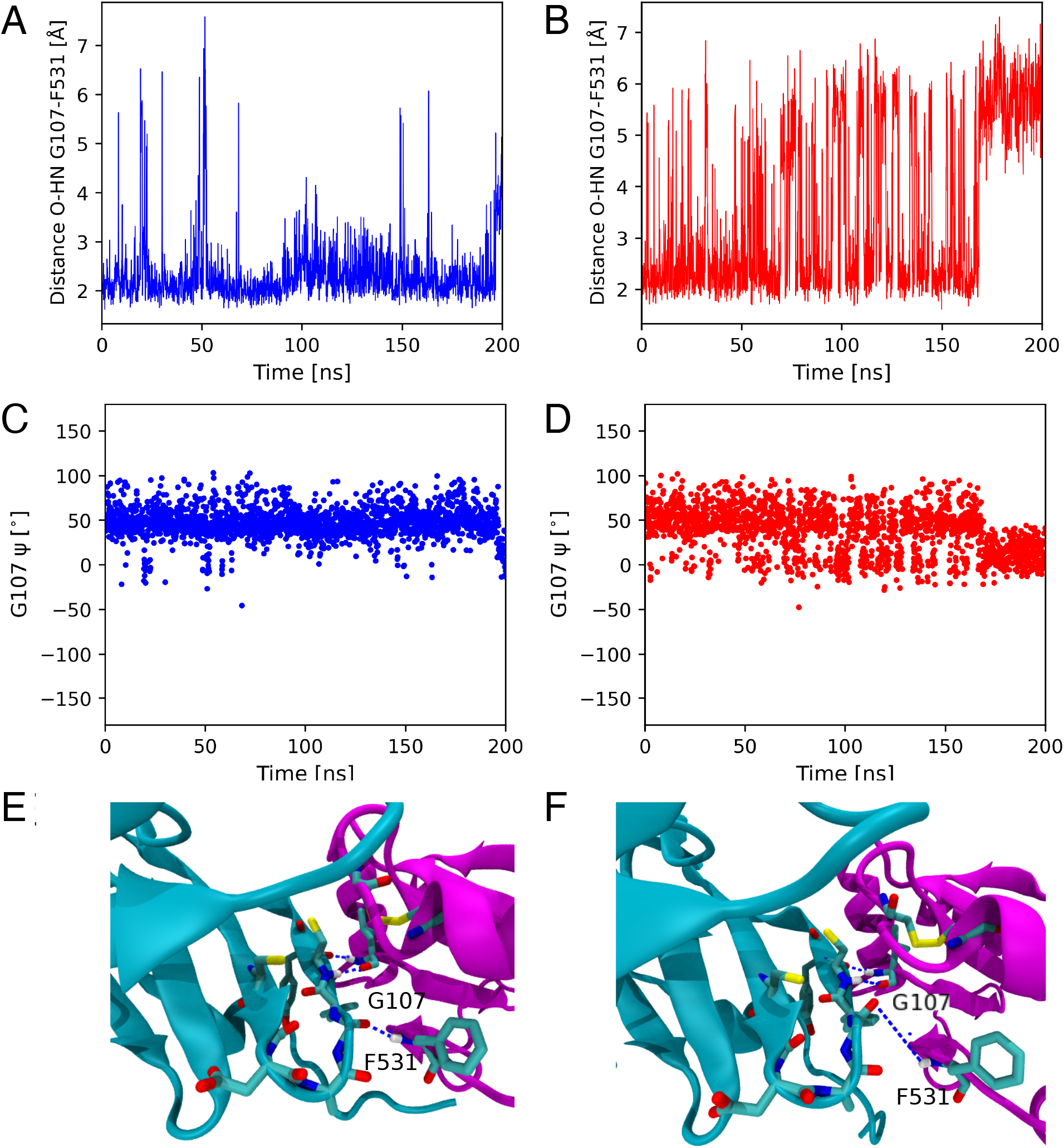
G107-F531 hydrogen bond is destablised when cDsbD active site is reduced. *ψ* dihedral angle of G107 in simulation of the (A) stable nDsbD_ox_-cDsbD_red_ and (B) the least stable nDsbD_red_-cDsbD_ox_ complexes. (C) and (D) G107-F531 hydrogen bond in simulations of the most stable, nDsbD_ox_-cDsbD_red_, and least stable, nDsbD_red_-cDsbD_ox_, complexes. Changes in *ψ* dihedral angle of G107 are correlated (A,B) with breaking the hydrogen bond (C,D). Simulation shows that the hydrogen bond nDsbD_red_-cDsbD_ox_ present in the starting configuration (C, at 0.2 ns) breaks. (D) Complex with dissociated hydrogen bond (at 200 ns).

The presence of thiols nDsbD_red_ rather than a disulfide likely changes the conformation of the active-site *β*-hairpin of nDsbD and de-stabilises the G107-F531 hydrogen bond. The cysteines in the active site of nDsbD in the complex adopt the same conformations as in nDsbD on its own (Fig. S7). The active site of nDsbD_red_ and in particular the C103 and C109 thiols provide a snugly fitting binding site for the cap loop of nDsbD and, thus nDsbD_red_ is less flexible^18^ than nDsbD_ox_. However, the larger size of the thiols in nDsbD_red_ as opposed to the disulfide bond in nDsbD_ox_ likely changes the *β*-hairpin conformation in the complex. In particular, the F108 *ϕ* (Fig. S8G,H,I) and *ψ* (Fig. S8J,K,L) angles show larger fluctuations in the simulation of nDsbD_red_. As the hydrogen bond breaks at around 170 ns, the F108 *ϕ* (Fig. S8H) and *ψ* (Fig. S8K) angles also change in nDsbD_red_. A similar albeit smaller shift is seen for nDsbD_ox_ at around 200 ns (Fig. S8G,J). These change in the backbone of the hairpin likely contribute to change in the orientation of the carbonyl of G107 and thus the stability of the G107-F531 hydrogen bond.

Running repeat simulations for 2 μs for nDsbD_ox_-cDsbD_red_ and nDsbD_red_-cDsbD_ox_ confirms the trend in stability of the inter-domain hydrogen bonds (Fig. S10). G107-F531 is more easily broken in simulation of the product complex cDsbD_ox_-nDsbD_red_ than in the simulation of the productive complex cDsbD_red_nDsbD_ox_, which is formed before of the electron transfer (Fig 1). In the repeat simulations of both complexes, the G107-F531 hydrogen bond breaks and reforms repeatedly which suggest that local equilibrium was established in both simulations, but the hydrogen bond is broken for longer periods of time in the simulation of the product complex. The bidentate C109 HN to L510 O and C109 O to L510 HN hydrogen bonds were stable in both repeat simulations in line with the initial simulations (Fig. S9).

The cap loop is flexible in the simulations of the cDsbD_red_nDsbD_ox_ and the cDsbD_ox_-nDsbD_red_ complex. Coupling between the active site and the cap loop in nDsbD on its own has been highlighted by NMR experiments and MD simulations.^18^ The cap loop shows considerable flexibility in both the stable and the less-stable complex (Fig. S11), while remaining in an open conformation. In both complexes, F70 seems to make larger excursions than Y71. We did not observe a clear coupling between active-site redox state and the cap-loop dynamics of the productive (cDsbD_red_-nDsbD_ox_) and product (cDsbD_ox_-nDsbD_red_) complexes.

### Investigating the full-length DsbD structure from AlphaFold in atomistic MD simulations

Building AlphaFold models of DsbD further enabled us to explore the interactions of the periplasmic domains in the context of constraints that might be imposed by the tethering of these domains to the transmembrane domain of DsbD. The helical secondary structure in the transmembrane domain closely matches the secondary structure previously predicted^33^ (Fig. 5A), the main differences appear to be the extent of TM3 (yellow, predicted HMMTOP 225-249). Searching for similar structures for the transmembrane domain using Foldseek,^34^ we find good agreement with the transmembrane domain of CcdA (PDB: 5vkv)^22^ (Fig. 5B). DALI^35^ aligns the two structures with a RMSD of 3.6 *°*A. The CcdA structure was determined by NMR and so it was not used to train AlphaFold^36^ and thus provides independent support for the predicted transmembrane structure. The conformation of helix 4 differs between CcdA and DsbD and DsbD features two additional transmembrane helices. The backbone of the C- and N-terminal domains of the AlphaFold model align closely with the X-ray structure of the covalent complex of cDsbD and nDsbD (CA RMSD 1.1 *°*A, for residues 12-122 and 432-540, 1VRS chains B and E). F70 and Y71 of the cap loop of nDsbD points away from the cysteines in the active site as in the 1VRS X-ray structure. Such a conformation with the two periplasmic bound to each other is an important intermediate in the functional cycle of DsbD and considering the close proximity of the two periplasmic domains such a conformation may be populated to some extent. It is important to point out that the most populated structure *in vitro* and *in vivo* will depend on the redox-state of the different domains.^23^ The catalytic cysteine pairs are modelled as oxidised by default by AlphaFold.. We expect conformation in which the two domains are dissociated, and not interacting with each other, to also play important roles since these conformations are critical for DsbD’s function as a redox hub in the periplasm.

**Figure 5:**
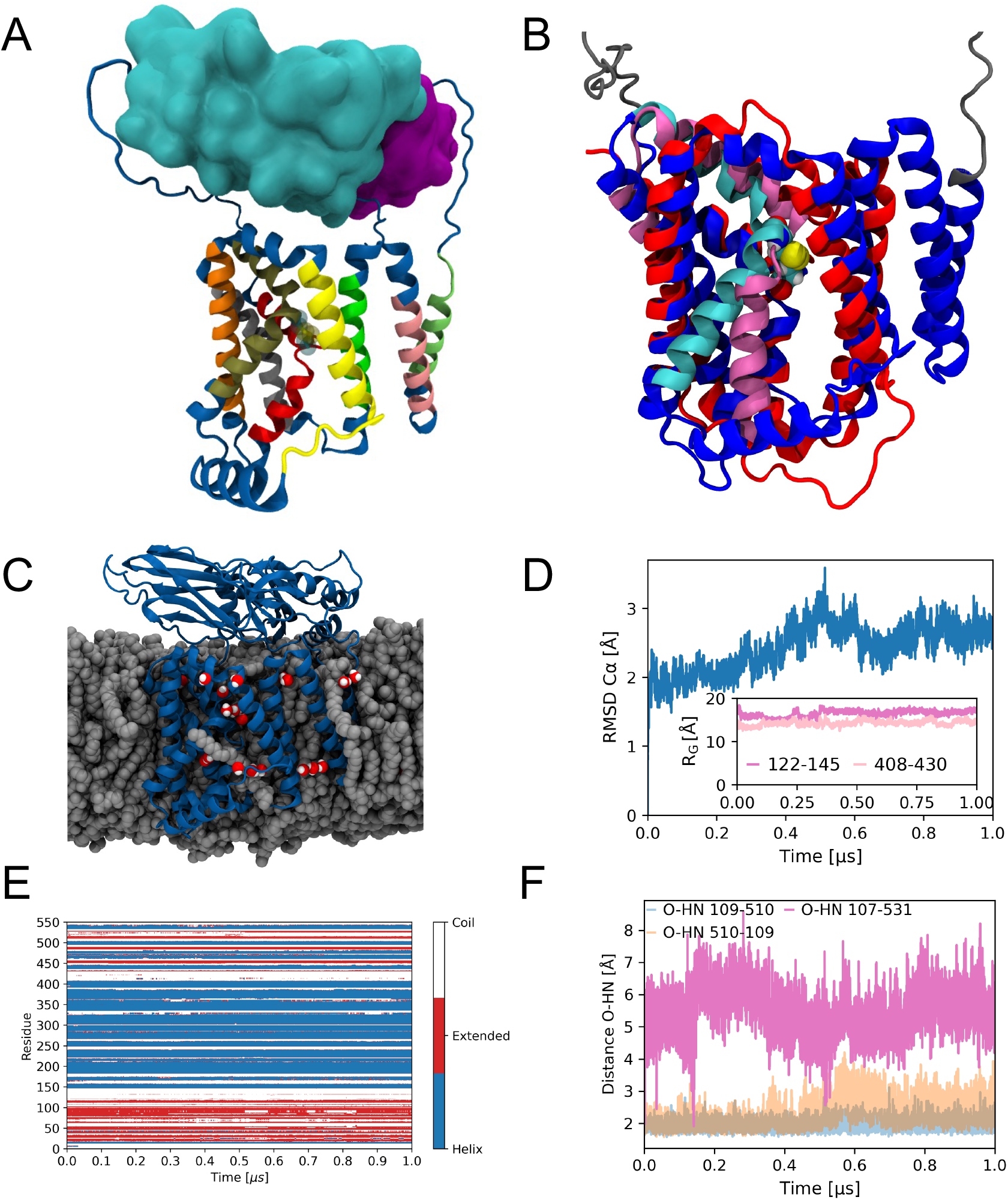
Full-length model of DsbD. (A) Comparison of predicted transmembrane helices with AlphaFold model. The colored segments are the helices predicted by Beckwith et al. nDsbD and cDsbD are shown in cyan and purple. (B) Transmembrane domain of DsbD AlphaFold model aligned with the NMR structure of CcdA. Transmembrane DsbD (145-408) in blue, linkers in gray. CcdA is shown in red. Helix 4 of DsbD and CcdA are highlighted in cyan and pink respectively. (C) Final structure from 1 μs of all-atom simulations with explicit solvent and membrane. Water is only shown for the vicinity of the transmembrane domain. (D) CA RMSD of the transmembrane domain (residues 145-408) compared to the equilibrated starting structure. Inset shows the extension of the linkers connecting nDsbD and cDsbD to the transmembrane domain. (E) Secondary structure over the course of the 1 μs simulation. (F) Hydrogen bonds between C109-G510 and G107-F531 at the interaction interface of nDsbD and cDsbD in the simulation

To better gauge what the model of full-length DsbD implies for its function we subjected it to atomistic molecular dynamics simulation with an explicit representation of a bacterial membrane (Fig. 6C). The transmembrane structure is maintained throughout 1 μs of atomistic molecular dynamics simulations. The CA RMSD (residues 145-408) to the AlphaFold model stays generally below 3 *°*A(Fig.5D). The linkers are highly dynamic as we would expect and it is reassuring that the state-of-the-art CHARMM36 simulation model^37^ captures the flexibility of these linkers. Secondary structure is preserved in all three domains (Fig.5E). Interestingly the helices in the transmembrane domain are maintained. Helices can break and re-form on the 100 ns time scale and consequently we can conclude that the transmembrane domain is in a favourable conformation in our model. The central space of the transmembrane domain of DsbD is partially hydrated,^38^ but no solvent filled cavity connecting the two different sides of the membrane is apparent (Fig.5C), which would be consistent with previous biochemical studies that argued against a continuous channel. ^33^ Interestingly, lipids protrude between the transmembrane helices and the extent to which this is relevant to the biological function of DsbD could be investigated in future studies (Fig.5C). In the AlphaFold structure the G107-F531 hydrogen bond between nDsbD and cDsbD is absent and this hydrogen bond does not form during 1 μs of simulations (Fig.5F). The bidentate hydrogen bonds between C109-L510 are mostly stable throughout the simulation, with the O L510 HN C109 distance increasing towards the end of the simulation, while HN L510 O C109 remain at a close distance throughout. nDsbD and cDsbD maintain a binding interface for the entirety of the simulations. This may force the linkers, connecting nDsbD and cDsbD (residues 122-145 and residues 408-430) to the transmembrane domain to remain in extended conformations throughout the simulations. The linkers are in coil conformations (Fig.5E) and the radius of gyration (*R*_*G*_) values computed for the two linkers (Fig.5D inset) are large considering a prototypical unfolded peptide of 24 amino acids for which we would expect a *R*_*G*_ value of 13 Å, *R*_*G*_ ∼ *R*_0_*Nν*, where *R*_0_ = 1.927 *°*A and *ν* = 0.598.^39^ In the absence of a model for full-length DsbD, the length of the linkers and to what extent they would be extended when the periplasmic domains interact has not been clear. The linkers are enriched in disorder promoting residues as reported by the CIDER webserver (http://pappulab.wustl.edu/CIDER/analysis/),^40^ with about 70% of linker residues classified as disorder promoting. The linker connecting the N-terminal domain and the transmembrane domain (residues 122-145) features five proline residues which favour extended structures. Once nDsbD-cDsbD contacts are broken, the linkers may enable the two domains to move away from each other, but likely the relatively limited linker lengths means that these domains are frequently close to each other thus increasing their effective concentrations as proposed previously.^23^ The periplasmic domains are in proximity to the membrane headgroup regions and thus any electrostatic differences between different redox-states are likely shielded by the charges of the lipid headgroups and the counter ions. Thus, small differences in interface conformations rather than differences in electrostatics may be essential in achieving the redox-state dependence of interaction affinities which is important in modulating the functioning of electron transfer by DsbD. ^18,23^ Overall, the full-length model and atomistic molecular dynamics of the full-length model of DsbD are in line with our conclusions from NMR relaxation dispersion experiments and molecular dynamics simulations of the nDsbD-cDsbD complexes as well as previous work, which suggested that small differences in the active-site conformations would determine the affinity of different nDsbD cDsbD redox pairs for each other.

**Figure 6:**
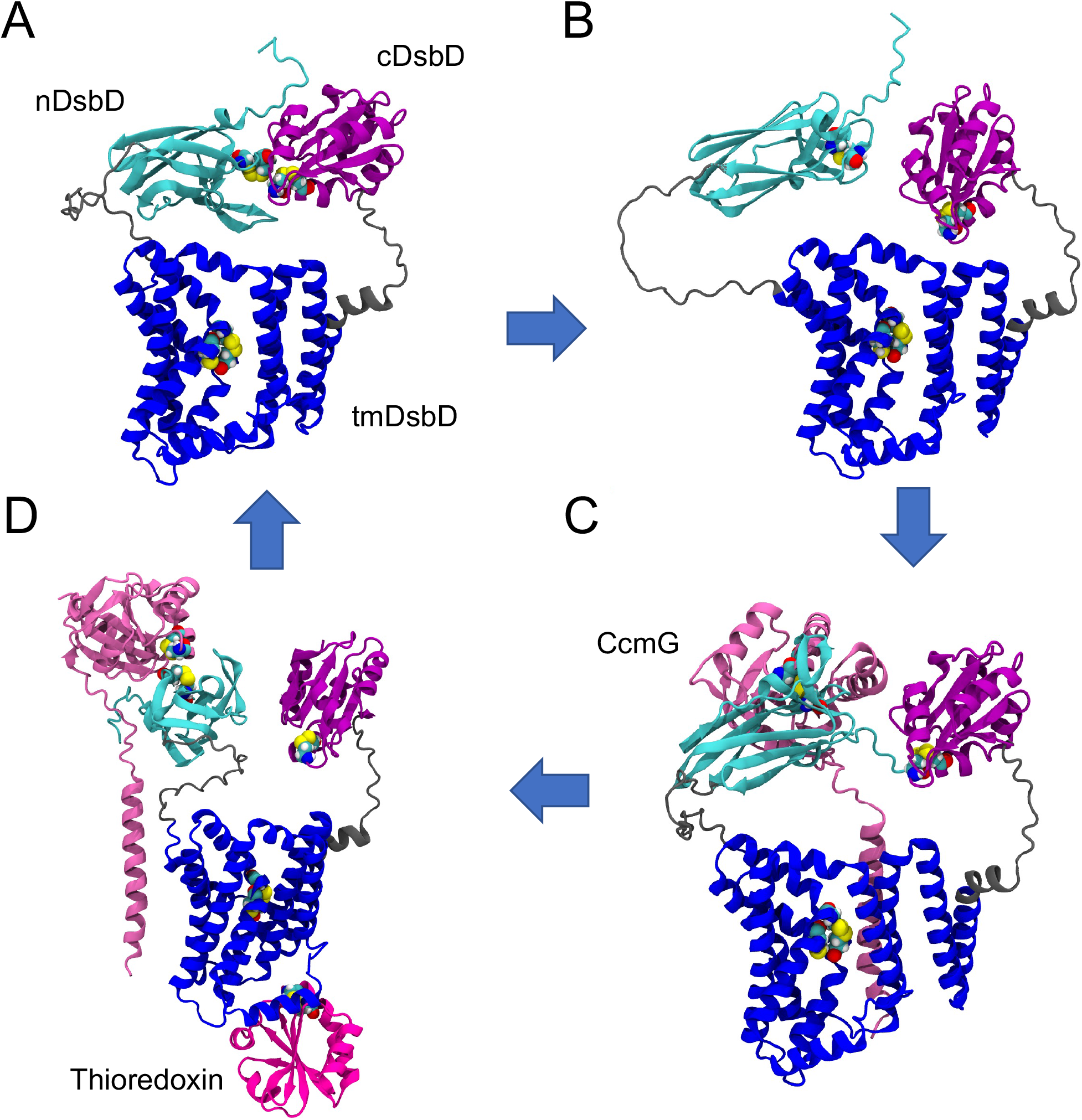
AlphaFold models of the functional cycle of DsbD. (A) Top-ranked full-length AlphaFold model of DsbD. Perpiplasmic domains nDsbD and cDsbD form a complex which enables electron transfer from cDsbD to nDsbD. (B) AlphaFold model (ranked 3rd) with nDsbD etached from cDsbD. Active site of cDsbD is orientated towards tmDsbD. (C) CcmG binding to nDsbD can be visualised by an AlphaFold model. Complex formation enables electron transfer to CcmG. (D) Thioredixin binds to re-supply tmDsbD with electrons. Complex of nDsbD and CcmG will dissociate and free up CcmG for its function in cytochrome c maturation. Electrons from tmDsbD are transferred in turn to cDsbD and new functional cycle starts.

### Modelling the functional cycle of DsbD with AlphaFold

With AlphaFold we can model further intermediates in the functional cycle of DsbD (Fig. 6A). Intriguingly, some models of full-length DsbD feature cDsbD and nDsbD dissociated from each other (Fig. 6B). In such conformations, nDsbD_red_ could bind to its cognate partners CcmG, DsbC, and DsbG and transfer electrons via a thiol-disulfide exchange reaction. To explore the coupling between the two periplasmic domains of DsbD we generated AlphaFold models in which either nDsbD or cDsbD were absent. In the absence of cDsbD, nDsbD does not pack closely to the transmembrane domain (Fig. S12A) and this structural plasticity may be important when nDsbD binds to its cognate interaction partners DsbC, CcmG, and DsbG in the bacterial periplasm, and transfers two electrons and two protons (reductant) to them. Interestingly, in the absence of nDsbD, cDsbD is re-orientated towards the interface of the transmembrane domain (Fig. S12B); in particular, C461 and C464 are brought into close proximity to the periplasmic face of the transmembrane domain. AlphaFold may be picking up co-evolution of the active site region of cDsbD and parts of the transmembrane domain. This closer proximity may be the first step leading to the close approach of the critical thiol groups of cDsbD and tmDsbD which is required for reduction of cDsbD. We are also able to generate a complex of DsbD with CcmG and thus model the next step in the functional cycle of DsbD (Fig. 6C). AlphaFold predicts that CcmG would bind to nDsbD in this complex, with cDsbD not interacting with nDsbD. Moreover, the transmembrane domain of CcmG is orientated towards the transmembrane domain of DsbD. Considering the positioning of DsbD in the membrane in atomistic MD (Fig. 5C), the transmembrane helix of CcmG would be orientated approximately parallel to the membrane normal. The AlphaFold model of CcmG bound to DsbD (Fig. 6C,D) and CcmG bound to a DsbD-thioredoxin complex reveal different contacts between the transmembrane domain of DsbD and CcmG and the interactions of the transmembrane domains may be fuzzy, which could help CcmG to dissociate once it has been reduced nDsbD (Fig. 6D). DsbD needs to be re-supplied with two electrons by thioredoxin for its next functional cycle (Fig. 1). Thioredoxin binds to the peripheral helix (residues 249-262). Its catalytic cysteines are orientated towards the transmembrane domain. The catalytic cysteines of thioredoxin are more than 20 *°*A away from the cysteines C163 and C285 in the transmembrane domain of DsbD, which suggests a conformational change is required for reductant transfer via a thiol-disulfide exchange reaction.

To understand possible large-scale conformational changes in the transmembrane domain which could play an important role in moving the electrons from thioredoxin across the membrane, we used an extension of AlphaFold designed to reveal alternative conformations.^41^ For the transmembrane domain we did not find large-scale conformational flexibility (Fig. S12C). This suggests that information about major conformational changes is absent in the transmembrane domain sequence and might be dependent on the presence and redox state of the globular domains of DsbD or the partners of this protein on either side of the cytoplasmic membrane.

### Correlating the effects of DsbD mutants on its function with structural features in AlphaFold models of DsbD

*In vivo* experiments of DsbD variants provide additional clues as to its mechanism in light of the AlphaFold models and support the location of the thioredoxin binding site predicted by AlphaFold. The covalent attachment of a reporter *c*-type heme from cytochrome *cd*_1_ is followed to asses the function of DsbD mutants *in vivo*.^24^ We investigated the function of the G206A, G155, and P284 variants, which were previously studied by Cho *et al* ^42^ and the G213A, P249A, G266A, and P333 variants that have never been explored previously. These experiments were conducted before the AlphaFold models were available and are therefore completely unbiased. Some mutations such as G206A and P284A do not affect DsbD function significantly (Fig. 7A). In previous experiments the P284A mutation reduced function considerably. ^42^ The inconsistency with the previous report could be explained by the difference in volumes of the cultures (we used 1 L of culture instead of 1 mL), and the expression time (24 h instead of 4 h); our experimental settings allow for much improved resolution in our functional assays. P284 is close to the catalytic cysteines in the transmembrane domain and models of DsbD suggest that P284 could help to bend transmembrane helix 4. The lack of strong phenotype may suggest that there are redundant interactions determining the shape of the transmembrane helix 4. By contrast, G266A and P333A mutations greatly reduce DsbD function (Fig. 7B). Considering that these two mutations are on the cytoplasmic side of DsbD, we wondered whether they might be close to a thioredoxin binding site (Fig. 1B). Indeed, both mutations are close to the thioredoxin binding sites predicted by AlphaFold (Fig. 6C). G266 is close to the N-terminus of transmembrane helix 4 and is nearby the end of the peripheral helix (residues 249-262) which may help to recognise thioredoxin, as shown in Fig. 7E. P333 is at end of a transmembrane helix and is also close to the thioredoxin binding interface. For the G255A and P333A variants of DsbD the amount of *c*-type cytochrome matured is comparable to that matured by C464A-DsbD, which is known to be entirely inactive; the small amount of signal detected for these variants is due to the fact that a small percentage of the apocytochrome gets immediately processed by the Ccm system directly after Sec translocation and, therefore, does not require DsbD for its maturation.^43^ G155A in transmembrane helix 1 reduces function to a lesser degree than P266 and P333, possibly as it is further away from a binding site of either cDsbD or thioredoxin.

**Figure 7:**
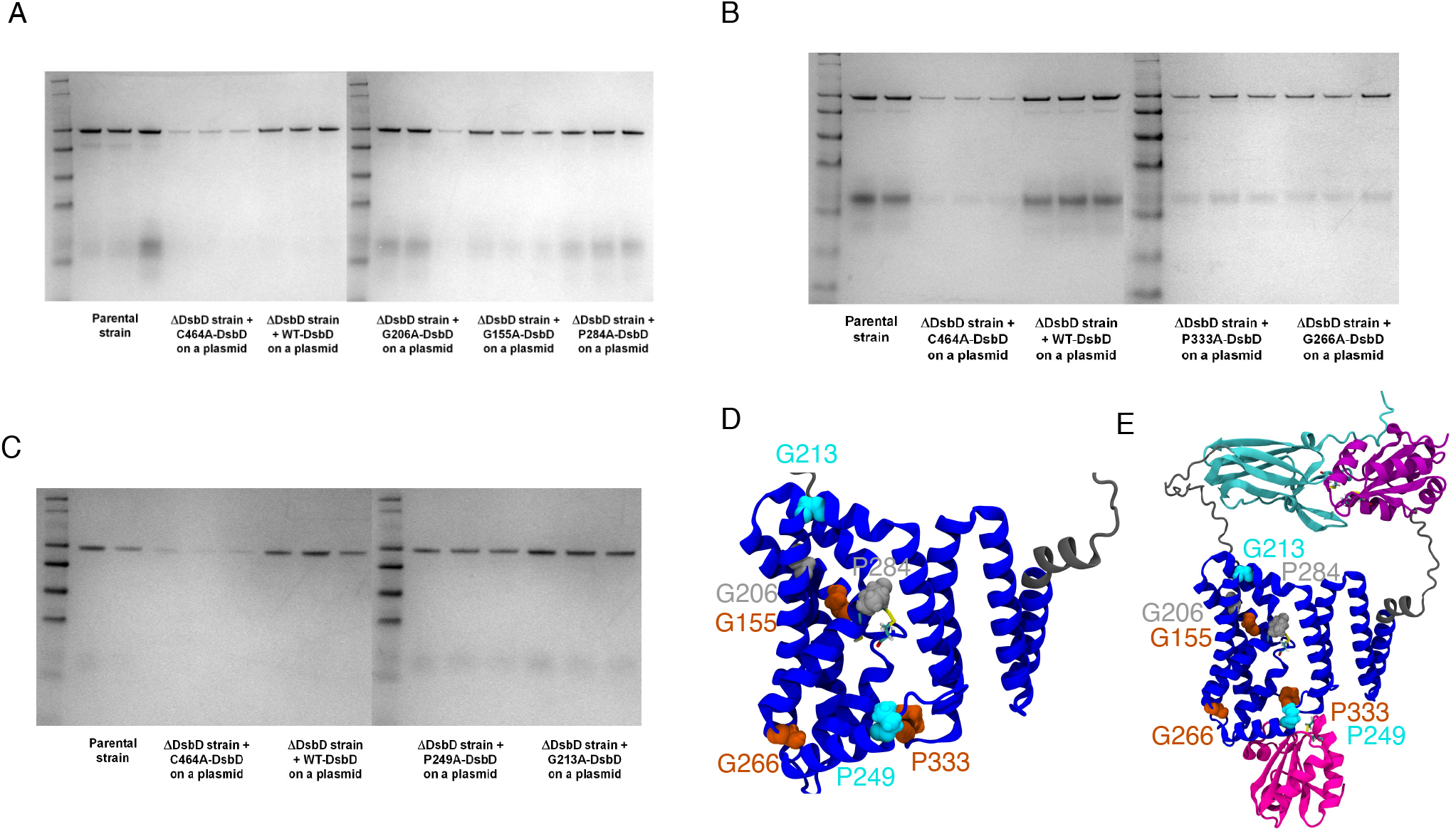
Effect of transmembrane mutations on the function of DsbD *in vivo* as reported by covalent heme *cd*_1_ attachment. *E. coli* was grown anaerobically. Triplicates are shown for each experimental condition. (A) Effect of G155A, G206, and P284A mutations on covalent heme attachment in the periplasm of *E. coli*. B) and C) effect of P249A and G213A as well as G266A respectively on covalent heme attachment *in vivo*. (D) Positions of the mutants in the transmembrane domain of DsbD from AlphaFold. Light blue are mutants that enhance DsbD function (G213 and P249). Orange are mutants that reduce DsbD function (G155, P333, G266). Mutants that do not change the activity are highlighted in light grey (P206 and P284). E) AlphaFold model of DsbD bound to thioredoxin. G266 is adjacent to the peripheral helix which may help to recognise thioredoxin. P249 is at the start of this peripheral helix. P333 is also close to the thioredoxin binding site

Surprisingly, two variants G213A and P249A, enhance the function of DsbD (Fig. 7C). G213 is towards the periplasmic end of the second transmembrane helix and while this mutation could affect local flexibility of the helix and local conformational dynamics, AlphaFold models do not suggest a clear explanation to the enhanced function of the P213A mutant. P249A might make the peripheral helix longer and this helix is right where thioredoxin binds. It is tempting to speculate that the P249A mutant helps thioredoxin binding by providing a larger recognition surface.

## Discussion

We combined NMR measurements, molecular dynamics and computational modelling, as well as *in vivo* experiments to elucidate the dynamic interactions of the different domains of DsbD. Macroscopic on and off rates were determined from the NMR relaxation dispersion experiments. The association rates for both C461A cDsbD and oxidised cDsbD_*o*_*x* with oxidised nDsbD_*o*_*x* were relatively fast, around 2×10^6^ M^*−*1^ s^*−*1^ and are approximately an order of magnitude faster than the rate constant for the reductant transfer reaction between reduced cDsbD and oxidised nDsbD.^21^ This might suggest that the reductant transfer between the two periplasmic domains is not limited by the kinetics of the complex formation^1^ but by the covalent chemistry of the thiol-disulfide exchange reaction. Oxidation of cDsbD has a large effect on *k*_off_ of the cDsbD-nDsbD complex. The cDsbD_ox_-nDsbD_ox_ complex dissociates with a *k*_off_ of about 2000 s^*−*1^, which is even faster than the dissociation rate of electron-transferring complexes.^9^ Note this effect on the dissociation rates of the domains on their own is likely even larger as the complex nDsbD_red_-cDsbD_ox_ of the products of the thiol-disulfide reaction interact only weakly.^23^

The intrinsic kinetics of the association and dissociation reaction of the periplasmic domains of the membrane protein DsbD are important for its function as an oxidoreductase and may be modulated by the linkers which connect cDsbD and nDsbD to the transmembrane domain of DsbD (Fig. 6B). Thus, cDsbD and nDsbD function at high local concentrations. Given the high local concentrations, nDsbD_ox_ is likely to encounter cDsbD when its cap loop is open as a result of the redox-state dependent dynamics of nDsbD.^18^ nDsbD_ox_ can then receive electrons from cDsbD_red_ via an intermolecular thiol-disulfide exchange reaction. Once the electrons are transferred the bulky thiols in the active site of cDsbD may disrupt the three key hydrogen bonds between cDsbD and nDsbD, speeding up the dissociation rate *k*_off_ of the complex. Our simulations show that the presence of thiols or disulfides in the active sites strongly modulates the stability of the inter-domain G107-F531 hydrogen bond, which is less-stable in the complex of the products cDsbD_ox_-nDsbD_red_. The cap loop of nDsbD_red_ is also less likely to open, as its active site is no longer structurally frustrated; ^18^ this means that once the product complex dissociates the complex is unlikely to form again. The twofold effect may be vital to prevent the thermodynamically feasible back reaction at high local concentrations of cDsbD and nDsbD.

The advent of AI-driven AlphaFold modelling provides for the first time a model of the transmembrane domain of DsbD and its likely interactions with the periplasmic domains and thioredoxin in the cytoplasm. The effect of mutants on covalent heme *cd*_1_ attachment in anaerobically grown *E. coli* is overall consistent with our structural modelling. For the related oxidoreductase CcdA an inverted-repeat topology has been proposed to underpin the movement of cysteines in the transmembrane domain to face either the cytoplasmic or periplasmic sides of the membrane in an elevator-type mechanism. ^22^ Our results suggest that the interactions of the periplasmic domains can be coupled to interactions of the periplasmic domains with the transmembrane domain (Fig. 6B and Fig. S12), which could trigger the elevator-type movement of the active site.

## Methods

The first sample, contained ^15^N labelled C461A-cDsbD with a final concentration of 0.55 mM. The final nDsbD concentration was estimated based on proton spectra (Fig. S1) to be about 25 fold less than the concentration of ^15^N labelled C461A-cDsbD. There was about 4 % D_2_Oin the sample. The pH was adjusted to 6.5.

In the second sample, the concentration of oxidised cDsbD was 0.53 mM. The final nDsbD concentration was estimated based on proton spectra (Fig. S1) to be about 17 fold less (Fig. S2) than the concentration of ^15^N labelled cDsbD_*ox*_. The pH was adjusted to 6.5 and 15 μL D_2_O was added. There was about 4 % D_2_O in the final sample volume.

### Spectrometers

NMR experiments discussed in this chapter were recorded on home-built 500 MHz, 750 MHz and 950 MHz spectrometers. The spectrometers are equipped with Oxford Instruments Co magnets, home-built triple-resonance probes with triple axis gradients. The spectrometers were controlled by GE/Omega data acquisition computers and software.

### NMR relaxation dispersion experiments

To study the association of ^15^N labelled C461A-cDsbD and unlabelled oxidised nDsbD, amide ^15^N relaxation dispersion experiments were collected at magnetic field strengths corresponding to proton resonance frequencies of 500 MHz and 950 MHz. To study the binding of ^15^N labelled oxidised cDsbD to unlabelled oxidised nDsbD, experiments were conducted at 500 MHz and 750 MHz. The pulse sequences introduced in the previous chapter were used. The length of the spin-echo train *T*_CPMG_ was 40 ms.

### Analysis of the relaxation dispersion experiments

Experiments were processed by standard methods using NMRpipe ^44^. The intensities of the amide resonances were followed in CCPNMR Analysis ^45^. The intensities were exported and converted to *R*_2,eff_ Relaxation dispersion curves were analysed using CATIA (Cpmg, Anti-trosy, and Trosy Intelligent Analysis, https://www.ucl.ac.uk/hansen-lab/catia/).^25–27^ CATIA fits the data by numerically integrating over a representation of the pulse sequence and is the most general way of analysing relaxation dispersion experiments. For both samples, a single global exchange process was assumed.

### Molecular dynamics simulations of cDsbD and nDsbD complexes and their formation

Simulations of complexes and the association reaction between cDsbD and nDsbD were setup based on the 1VRS B crystal structure.^21^ The two domains were then solvated in a rhombic dodecahedral box. The minimum distance between the protein and the box edges was 1.5 nm. The D455 side-chain is protonated in the complex of cDsbD and nDsbD and a proton was consequently added to this side-chain in the setup of the MD simulations.^16^ Histidine side-chains were assumed to be positively charged in all simulation setups. The system was energy minimised.

MD simulations were run in GROMACS 4.5 ^46,47^ using the CHARMM 22 potential energy function with the CMAP correction ^48^ following the approach outlined by Bjelkmar *et al*. The lengths of bonds involving hydrogen atoms was constrained using the PLINCs algorithm.^49^ The equations of motion were integrated with a 2 fs time step. To equilibrate the simulation system we run a 4 ns MD simulation with position restraints with harmonic force constants of 1000 kJ*/*mol^2^ on the protein heavy atoms before the production runs.. Electrostatics was treated using the Particle Mesh Ewald approach with a short range cutoff of 12 *°*A. The Lennard-Jones potential was switched to zero between 10 *°*A and 12 *°*A. The neighbour list was updated every five integration steps using a cut-off radius of 12 *°*A. The length of covalent bonds to hydrogen atoms was constrained using the PLINCs algorithm ^49^. An integration time step of 2 fs was used.

For simulations of the cDsbD- nDsbD complexes the Xray-structure was prepared in the respective redox-states. We simulated the cDsbD_red_-nDsbD_ox_ (complex before electron transfer) and cDsbD_ox_-nDsbD_red_ (complex after electron transfer). Simulations were run for 200 ns each. Repeat simulations were run with the CHARMM36 force field^37^ for both complexes for 2 μs with GROMACS/2021.^50^ The D455 side-chain was de-protonated in the repeat simulations.

For simulations of the association the 1VRS B crystal structure was used to orientate nDsbD and cDsbD. The 1L6P and 3PFU crystal structures ^23,51^ were aligned to the 1VRS B structure and subsequently translated, using MDAnalysis ^52^ and were separated by 50.The D455 side-chain is protonated in the complex of cDsbD and nDsbD and a proton was consequently added to this side-chain in the setup of the MD simulations. Ions were added to equilibrate the charge in the simulation system. Na^+^ and Cl^−^ ions up to a concentration of 100 mM were added to the first run. In a second run, no ions beyond what was needed to neutralise the system were added.

### AlphaFold modelling of DsbD and its complexes with thioredoxin

The full-length structure was generated with AlphaFold through ColabFold^2,53^ (https://github.com/sokrypton/ColabFold). The signal peptide, which is cleaved in cells, was excluded from the model generation. The model generation resulted in a full-length model with the functionally important cysteines in N- and C-terminal as well as the transmembrane domains of DsbD oxidized to disulfide bonds. Models without cDsbD (residues 408-546) and nDsbD (residues 1-144) and of complexes of DsbD with thioredoxin, and thioredoxin and CcmG were also generated with ColabFold.^2,53^ Possible alternative structures of the transmembrane domain were generated by the protocol of del Alamo *et al*. (https://github.com/delalamo/af2_conformations).^41^

### Molecular dynamics simulations of the full-length DsbD from AlphaFold

The full-length AlphaFold structure was inserted into a model bacterial bilayer with 455 POPE 130, POPG, and 65 CDL2 molecules using CHARMM-GUI.^54,55^ The membrane composition followed recent work Corey *et al*.^56^ Coarse-grained Martini simulations were used to equilibrate the full-length AlphaFold structures in the membrane. With the CG2AT approach (https://github.com/owenvickery/cg2at) of Stansfeld *et al*, ^57^ the AlphaFold full-length structure was fitted to the coarse-grained structure and the coarse-grained system was back-mapped into a fully atomistic representation. The final atomistic simulation system consisted of 402 000 atoms. In atomistic simulation of the protein membrane system, we used the CHARMM36^37^ force field. Coarse-grained and atomistic simulations were run with GROMACS/2021.^50^ Secondary structure content was followed by DSSP^58^ as implemented in mdtraj.^59^

### Analysis of MD simulations

Molecular structures and MD trajectories were visualized using VMD. ^60^ MD simulation trajectories were analyzed using the MDAnalysis Python package. ^52^

### *In vivo* cytochrome *cd*_1_ attachment assay of DsbD mutants

To assess the function of DsbD mutants *in vivo*, cytochrome *cd*1 from *Paracoccus denitrificans* was expressed from a plasmid, using a strategy previously described.^24^ *E. coli* were grown anaerobically in 1 L cultures. The native *Ccm* system is expressed when *E. coli* is grown anerobically.

As a positive control we used the partental *E. coli* strain with the same plasmid. As a negative control we employed an *E. coli* strain with catalytically inactive DsbD with the C464A mutation^61^ and the plasmid coding for cytochrome *cd*1 from *Paracoccus denitrificans*. We harvested the cultures after 24 h of growth and analysed periplasmic extracts with SDS-PAGE. We used the method of Goodhew^62^ to stain proteins with covalently attached heme and used densitometry to compare the heme-stained bands. The intensity of the bands of the parental strain with the cytochrome *cd*1 plasmid was taken as reference for comparisons.

## Acknowledgement

L.S.S. acknowledges the Biotechnology and Biological Sciences Research Council (BBSRC) for a graduate studentship (BB/F01709X/1). D.A.I.M. acknowledges funding from the National Institute of Allergy and Infectious Diseases of the National Institutes of Health (Award Number R01 AI158753) and The University of Texas at Austin Faculty Support Funds; the content is solely the responsibility of the authors and does not necessarily represent the official views of the National Institutes of Health.. M.S.P.S. acknowledges funding from the Wellcome Trust (grant number 208361/Z/17/Z) and the BBSRC (grant number BB/R00126X/1). C.R. acknowledges funding from the Wellcome Trust (grant numbers 079440, 092532). L.S.S. acknowledges support by ReALity (Resilience, Adaptation and Longevity), M^3^ODEL and Forschungsinitiative des Landes Rheinland-Pfalz. The authors gratefully acknowledge the computing time granted on the supercomputer Mogon (hpc.uni-mainz.de). We thank Prof. D. Flemming Hansen and Dr. Martin Vögele for helpful discussions, and Nick Soffe for help with the NMR pulse sequence programming.

## Supporting information

### 1D ^1^H NMR experiments

We used 1D ^1^H NMR experiments to track the relative concentrations of labelled cDsbD, C461A cDsbD (Fig. S1) or oxidised cDsbD (Fig. S2), and unlabelled nDsbD.

**Figure S1:**
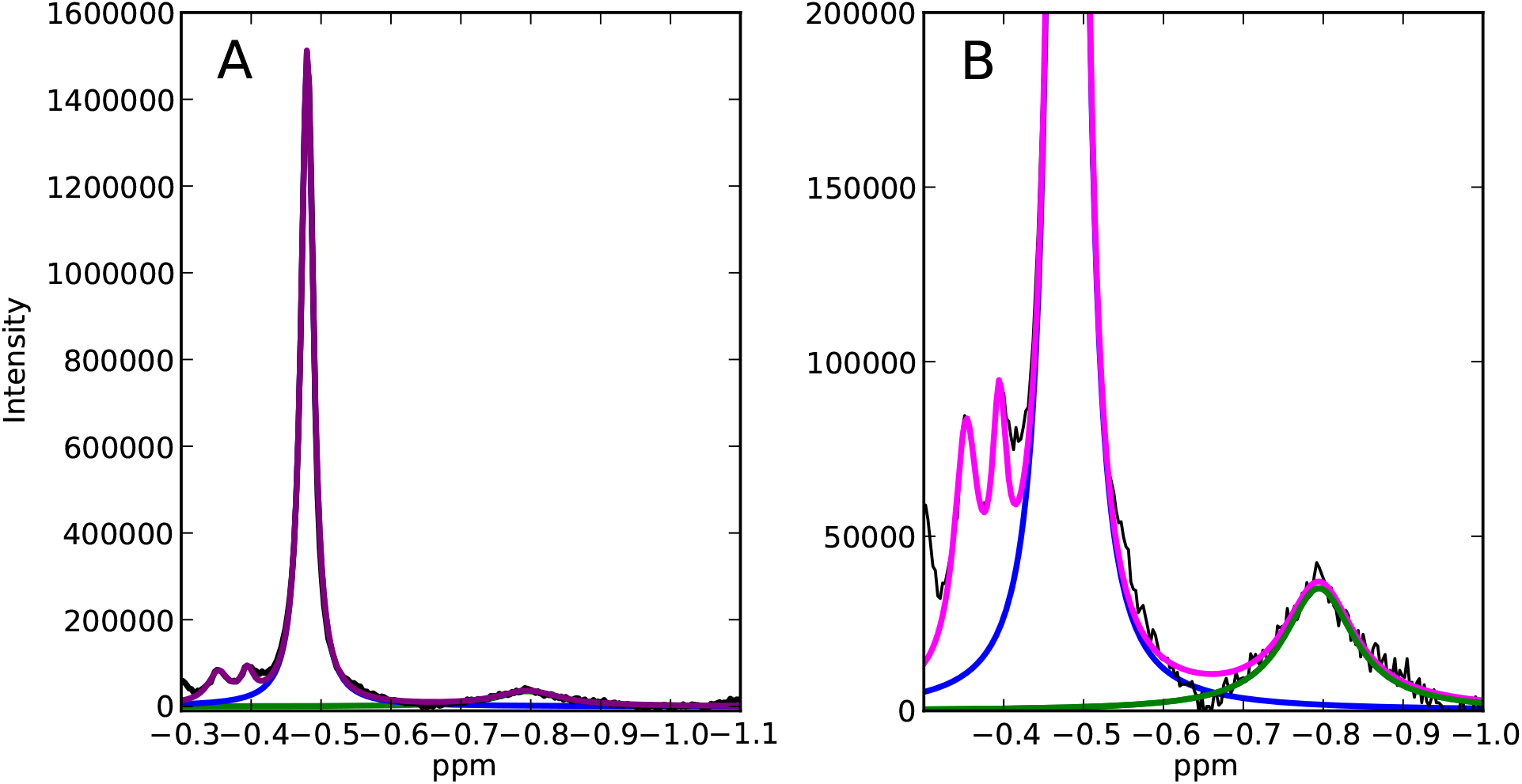
1D ^1^H NMR spectra tracking the relative concentrations of oxidised nDsbD and C461A cDsbD.

### NMR relaxation dispersion curves of ^15^N-C461A cDsbD in presence of oxidised nDsbD

**Figure S2:**
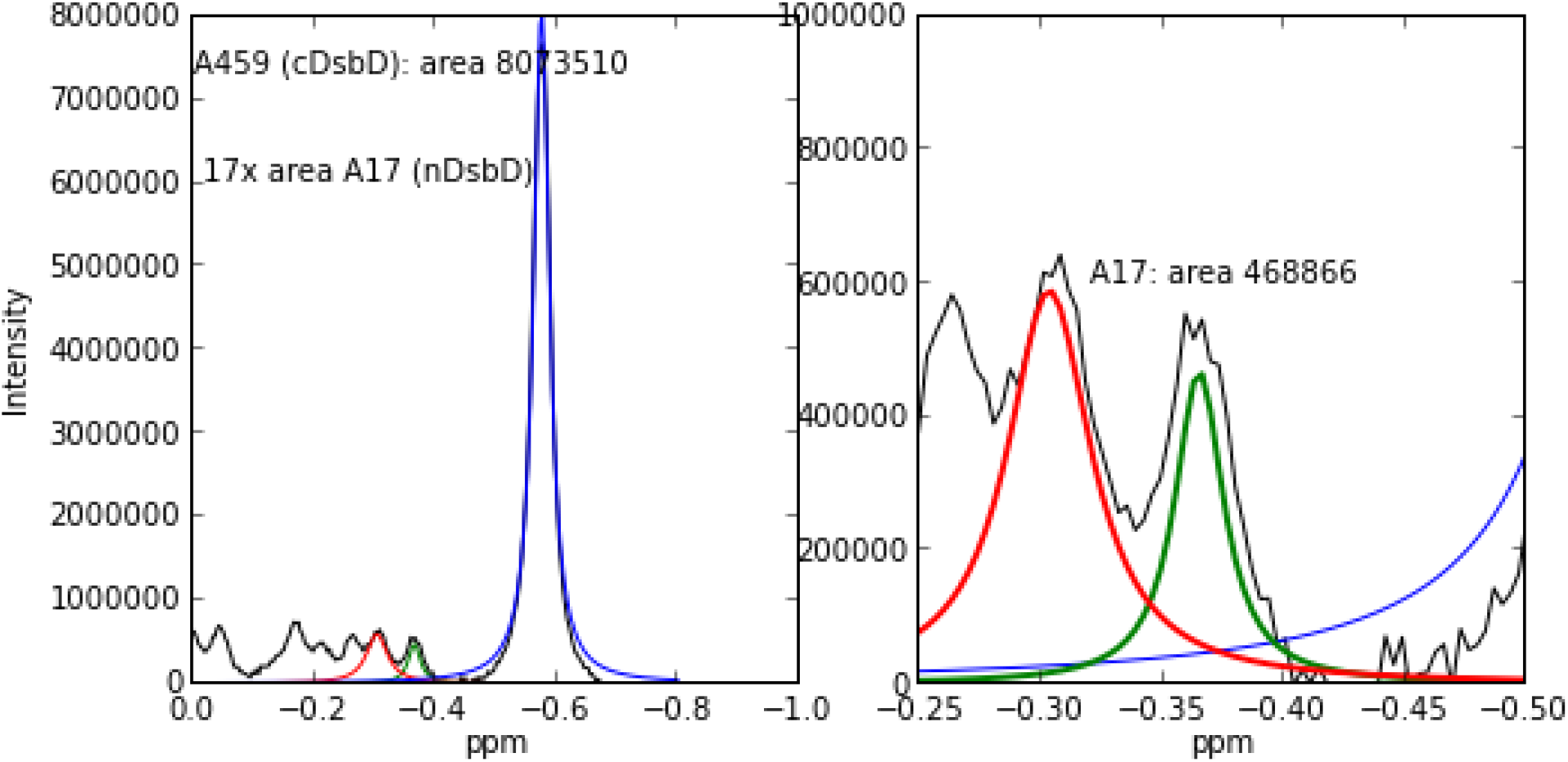
1D ^1^H NMR spectra tracking the relative concentrations of oxidised nDsbD and oxidised cDsbD.

**Figure S3:**
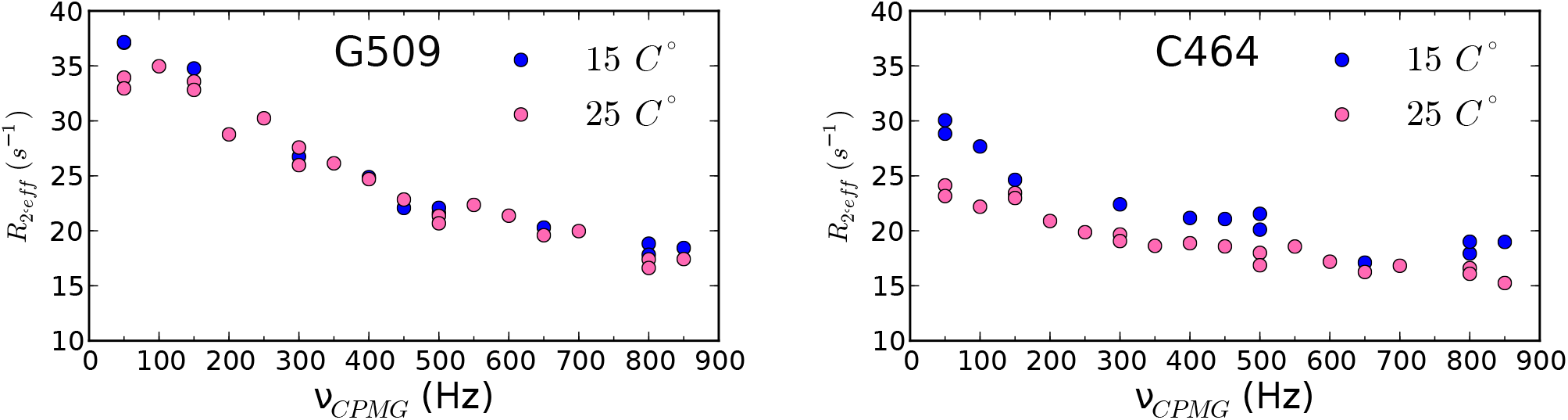
Relaxation dispersion experiments at 25 °C and 15 °C of ^15^N cDsbD_ox_ interacting substoichmetric amount of unlabelled nDsbD_ox_

**Figure S4:**
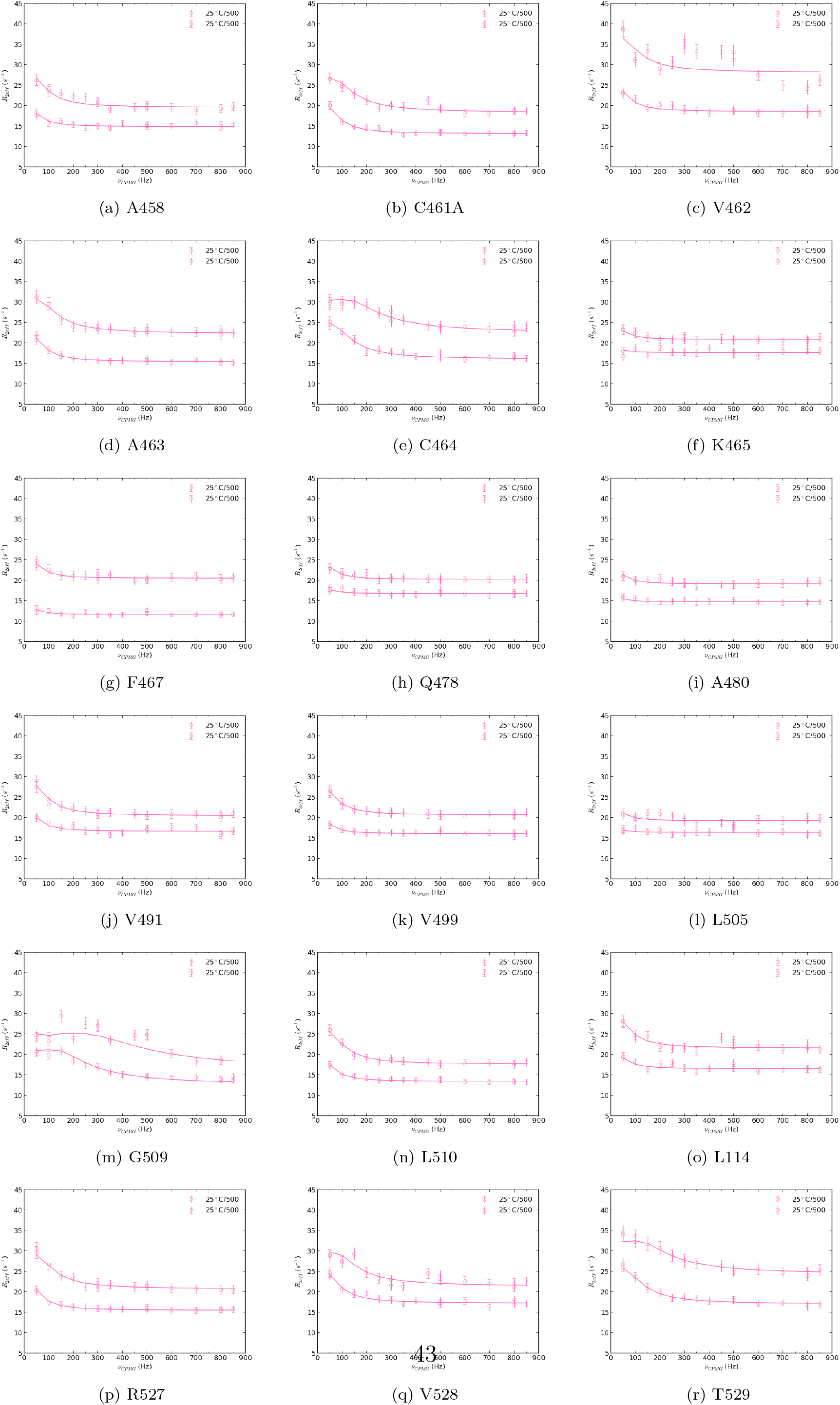
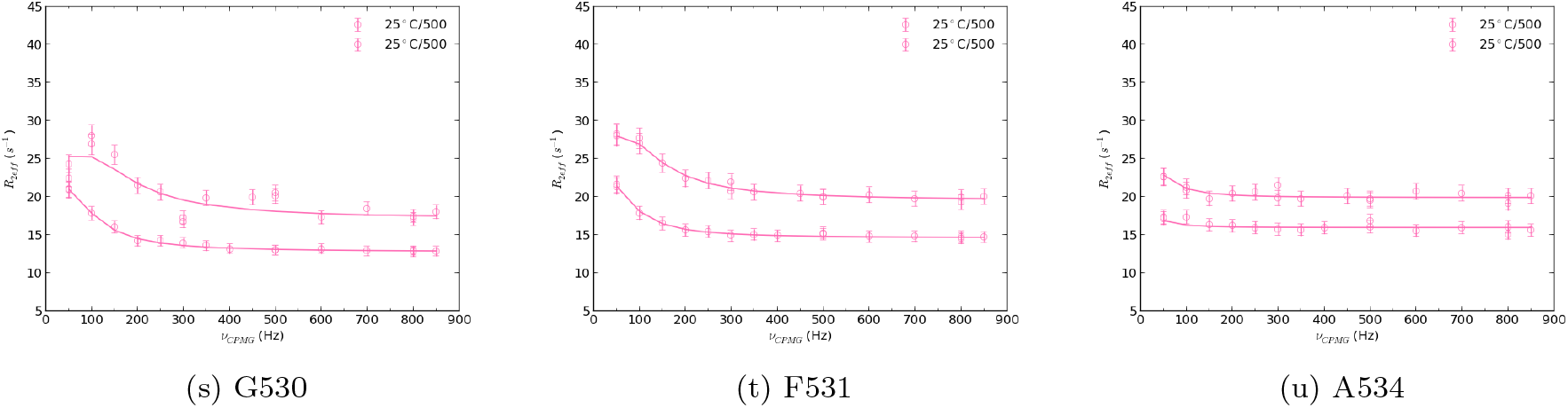
NMR relaxation dispersion experiments ^15^N-C461A cDsbD in presence of sub stochiometric oxidised nDsbD

### NMR relaxation dispersion curves of oxidised ^15^N cDsbD in presence of oxidised nDsbD

**Figure S5:**
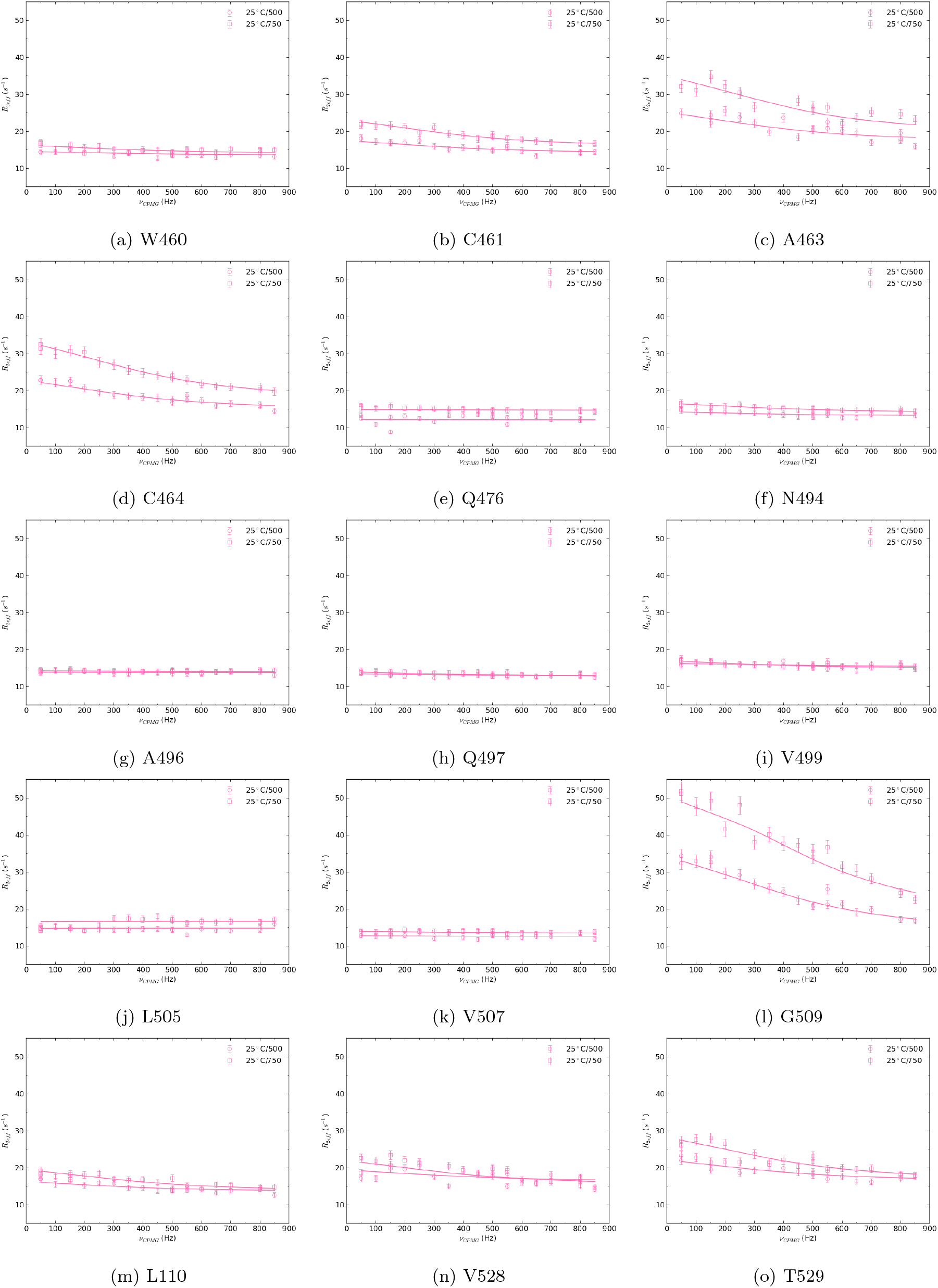
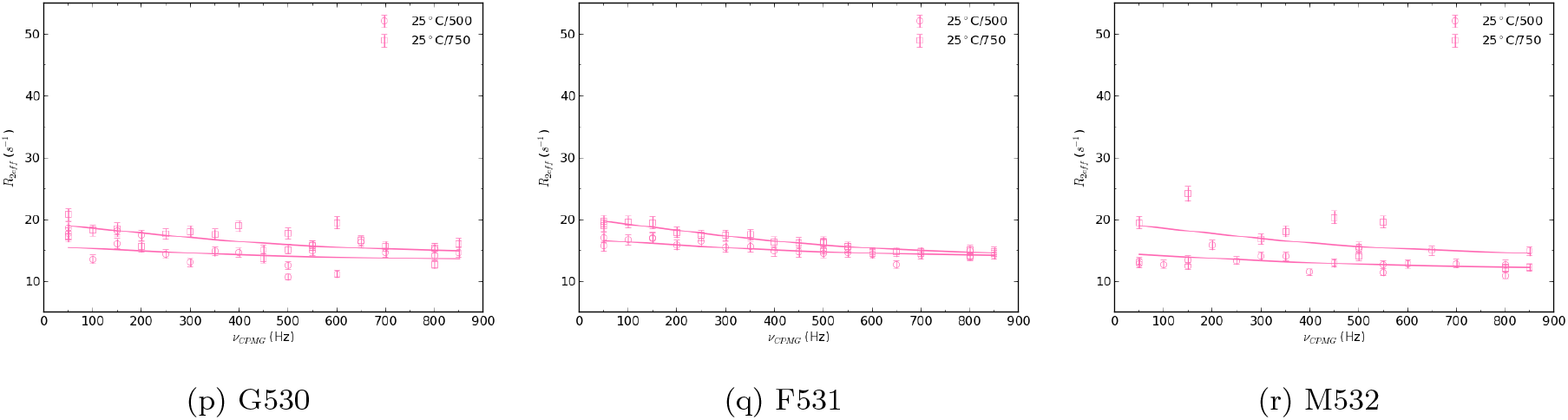
NMR relaxation dispersion experiments oxidised ^15^cDsbD in presence of substochiometric oxidised nDsbD

### Computational analysis of the association of cDsbD and nDsbD

#### Molecular dynamics simulations of the association of cDsbD and nDsbD

To better understand the association of cDsbD_red_ and nDsbD_ox_, we employed atomistic molecular dynamics (MD) simulations, electrostatic calculations and modelling of the association rate, which suggests that the association is not strongly enhanced by electrostatic interactions. We started MD simulations with the C- and N-terminal domains separated by about 50 *°*A. In the first simulation with 150 mM NaCl, the two domains associate quickly (Fig. 4A). While the C- and N-terminal domains are in contact at the cognate interaction interface (Fig.S6C), the cap loop remains closed and the native complex is not formed. The complex is stabilised by up to three inter-domain hydrogen bonds (Fig. 4C). The main hydrogen bonds are between HN C109 and O G509, T529 and E69, F531 and E69, as well as between R8 and V507. E.g., a long-lasting hydrogen bond was formed between the amine of C109 of nDsbD and the backbone carbonyl oxygen of G509 (Fig.S6D), right next to where L510 forms hydrogen bond with C109 in the X-ray structure of cDsbD-nDsbD. However none of the three native hydrogen bonds (O C109 and HN L510, O L510 and HN C109, O G107 and HN F531) were present in the 200 ns trajectory. ^21^ This complex might constitute a native-like encounter complex. In the second simulation run, no salt was added and we only added ions to neutralize the net charge of the protein domains. This setup will accentuate electrostatic interactions between the two domains and can possibly reveal electrostatic steering or long-range polar attraction.^13^ In the initial part of the trajectory cDsbD_red_ and nDsbD_ox_ appear to repel each other. Nonetheless an encounter complex is eventually formed after about SI50ns of simulation. As in the first simulation, the cap loop did not open and the native-complex like in the 1VRS X-ray structure was not accessed Interestingly, this complex behaves very differently from the one in the presence of 100 mM NaCl. The distance between the two domains fluctuates even after the initial formation of the complex as shown by Fig. S6E. The complex is very dynamic. Closely packed configurations, such as highlighted in Fig S6F exchange with much looser arrangements (Fig S6G). In such conformations the inter-facial region between the two domains can be largely solvated. Moreover, visual inspection of the trajectory showed that different parts of the two proteins interact with each other over time. For example, at 78 ns the C-terminus of cDsbD is in contact with the segment preceding the cap loop of nDsbD (Fig. S6H), with the side-chain of Q521 and the carbonyl oxygen of G63 forming a hydrogen bond transiently. About 10 ns later, the C-terminus of cDsbD has moved along the surface of nDsbD to interact with its C-terminal region (Fig. S6I).

**Figure S6:**
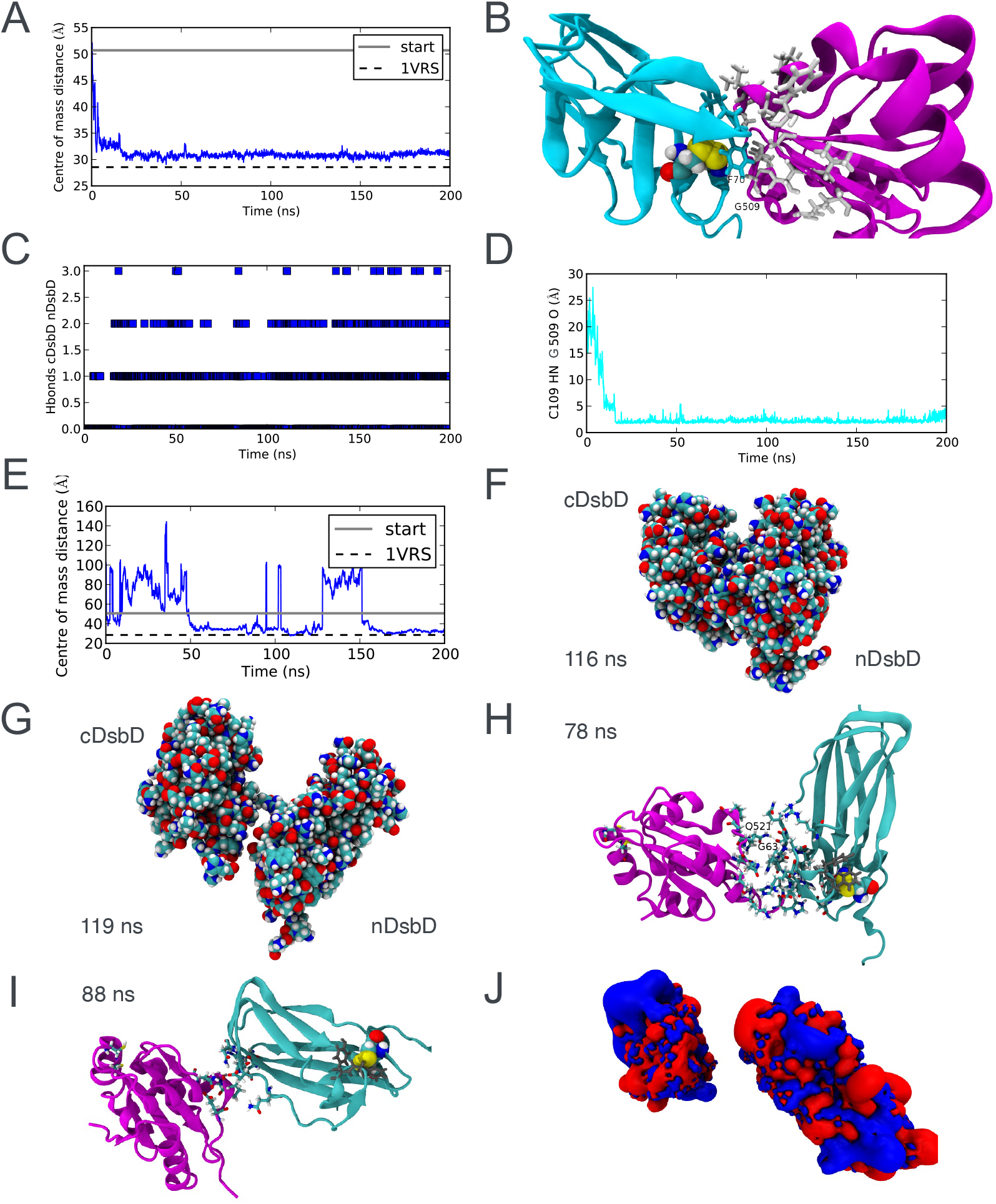
Modelling of the association of cDsbD_red_ and cDsbD_ox_. (A) Distance between cDsbD and nDsbD in simulation with salt. (B) A trapped state involving the cognate interaction interface but with a closed cap loop. cDsbD with amide nitrogen chemical shift changes of more than 1 ppm are highlighted in white. (C) Number of inter-domain hydrogen bonds. (D) C109 NH - G509 O hydrogen bond in the simulation. (E) Distance between cDsbD and nDsbD in simulation with no salt. (F,G) Closely packed (F) and loosely packed conformations exchange (H,I) Unspecific interactions in the simulations of cDsbD and nDsbD in the absence of additional salt. (I) Close, non-native interactions between the C-terminus of cDsbD and the region preceding the cap loop of nDsbD at 78 ns of simulation time. (I) At 88 ns, the C-termini of both domains interact. The cap loop (in grey) remained packed onto the disulfide bond of nDsbD (vdW representation). (J) Electrostatic potential surfaces are shown for cDsbD (to the left) and nDsbD (to right). The iso-contour surfaces are shown at +ve and -ve 2 KTe^−^ in blue and red, where *kT* is the thermal energy at 25 °C and *e* is the charge of an electron.

#### Electrostatics calculations

The Poisson Boltzmann equation was solved using APBS.^63^ The electrostatic potential surface was calculated for reduced cDsbD and oxidised nDsbD with the centre of masses separated by about 50. The charges were assigned using pdb2pqr.^64^ The D455 side-chain is protonated in the complex of cDsbD and nDsbD^17^ and a proton was consequently added to this side-chain. Histidine side-chains were assumed to be positively charged. The calculation was run at zero ionic strength to reveal potential complementary interactions surfaces under conditions similar to the low salt conditions employed in the NMR experiments.

Electrostatics calculations show that there is a large negatively charged patch around the cap loop of nDsbD, but there is not a clearly defined complementary positively charged surface on cDsbD, which would accelerate the association by electrostatically steering the two domains towards complex formation (Fig. S6J) Importantly, the full-length model of DsbD implies that the cDsbD-nDsbD complex is formed close to the surface of the membrane. Charged lipids in the membrane would also argue against a functionally role of electrostatic modulation of *k*_on_.

#### Transition Complex Theory

Transition Complex Theory (TCT)^30,31^ predicts an association rate *k*_on_ of 3.65 × 10^5^ M^*−*1^ s^*−*1^ roughly an order magnitude slower than the experimentlly determined rate coefficent. Note that TCT, which greatly simplifies the calculation of the association rate, can only provide approximate orders of magnitude estimates. Importantly, electrostatics does not enhance the interaction according to TCT. Considering these results, redox-dependent changes in the electrostatic surface of cDsbD are unlikely to affect *k*_on_, consistent with experiments.

### Additional analysis of the MD simulations of the cDsbD-nDsbD complexes

#### Cysteine conformation in the active site of nDsbD

**Figure S7:**
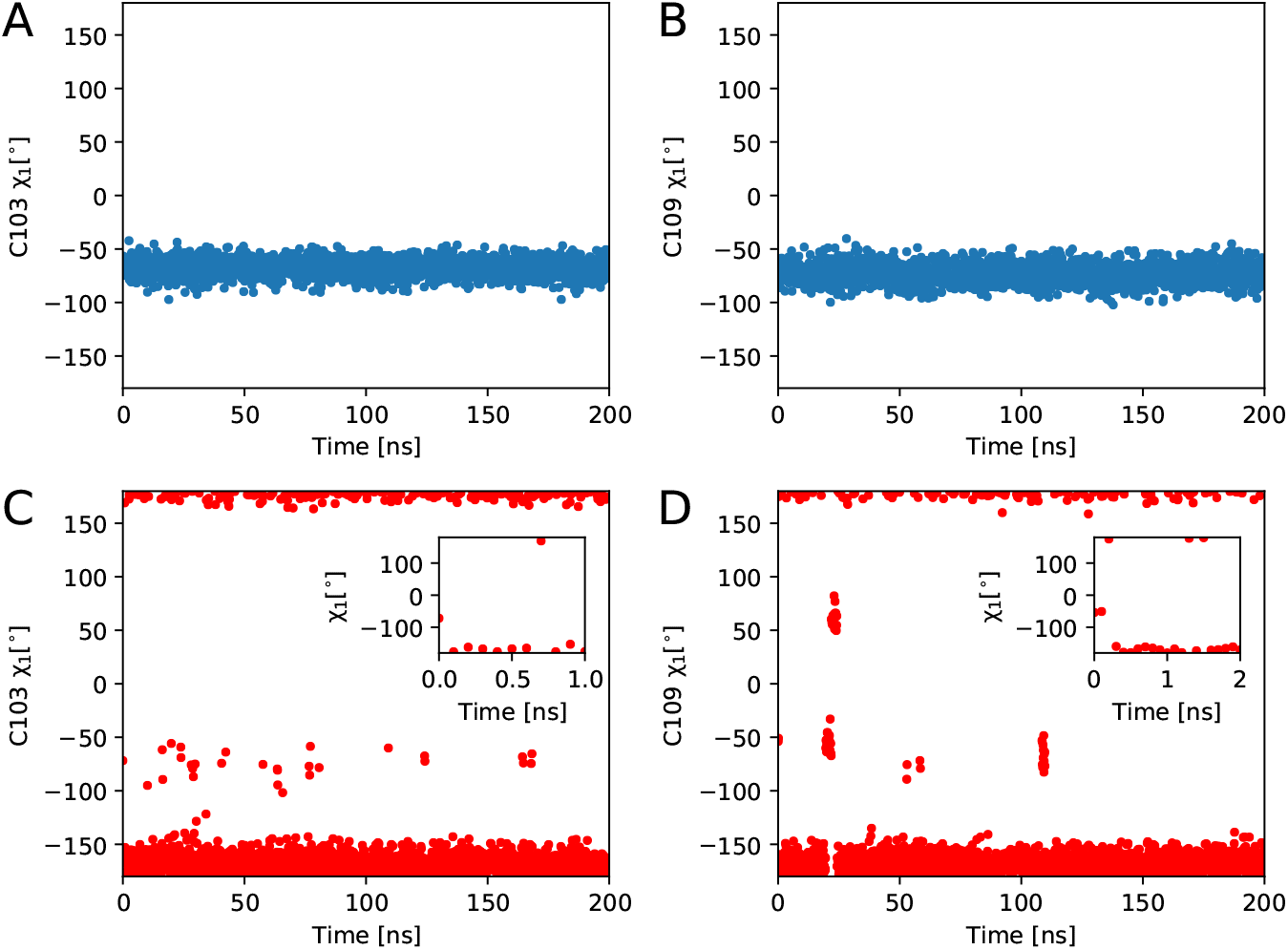
nDsbD active-site cysteines in stable cDsbD_red_-nDsbD_ox_ (A,B) and less stable (C,D) cDsbD_ox_-nDsbD_red_ complexes adopt gauche- and trans conformations as in nDsbD on its own.^18^ Inset in (C,D) shows initial relaxation of cysteines in reduced nDsbD in the less-stable complex.

#### Backbone dihedral angles in simulations of cDsbD-nDsbD complexes

The presence or absence of a disulfide bond leads to some differences in the conformation of the active site *β* hairpin as tracked by differences in the backbone dihedral angles (Fig. S8).

The C109 O L510 N and C109 O and L510 hydrogen bonds were maintained in the simulations (Fig. S9).

**Figure S8:**
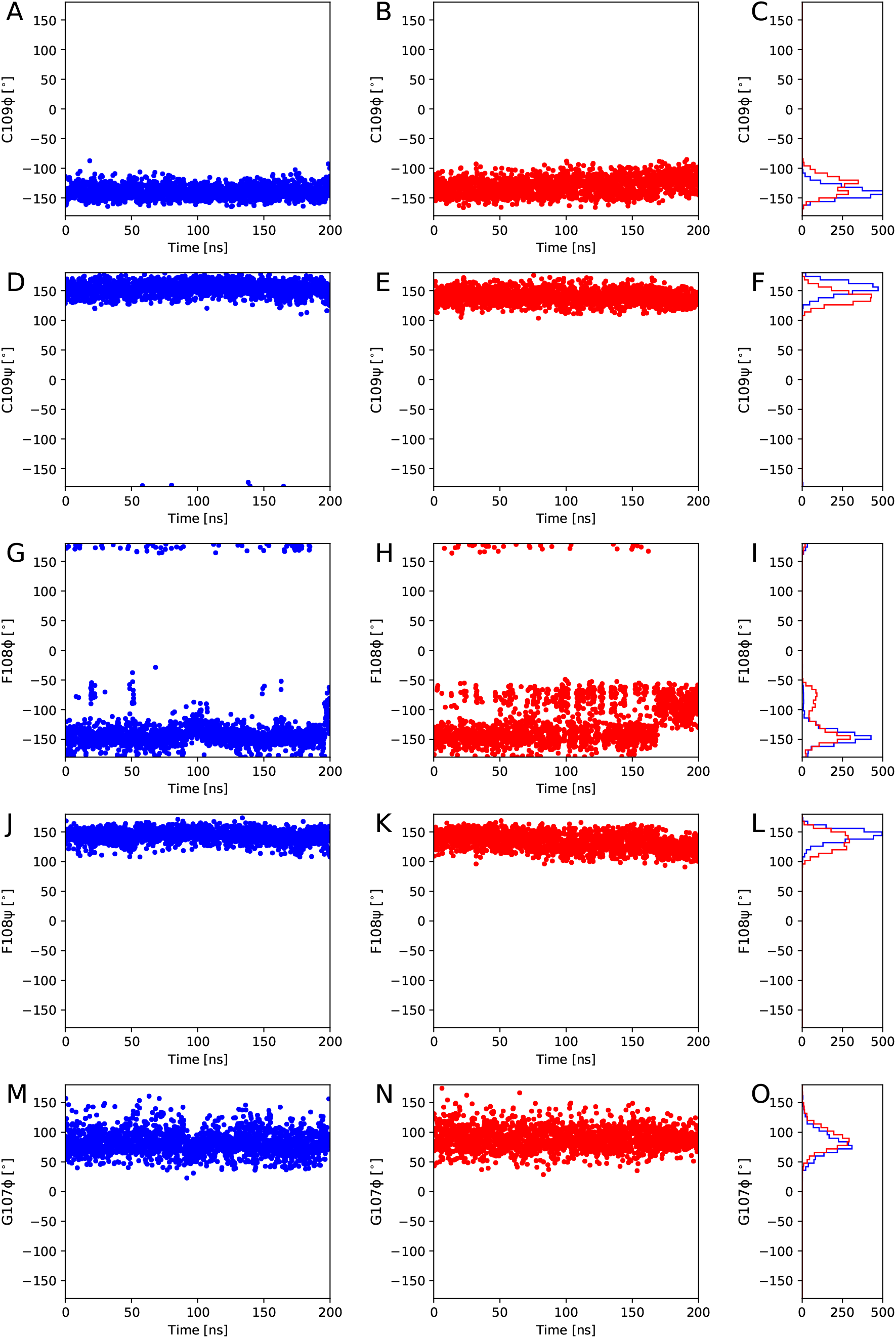
Differences in backbone dihedral angles in the simulation of the (A) stable cDsbD_red_-nDsbD_ox_ and (B) less stable cDsbD_ox_-nDsbD_red_ complexes.

#### Inter-domain hydrogen bonds in simulations of cDsbD-nDsbD complexes

**Figure S9:**
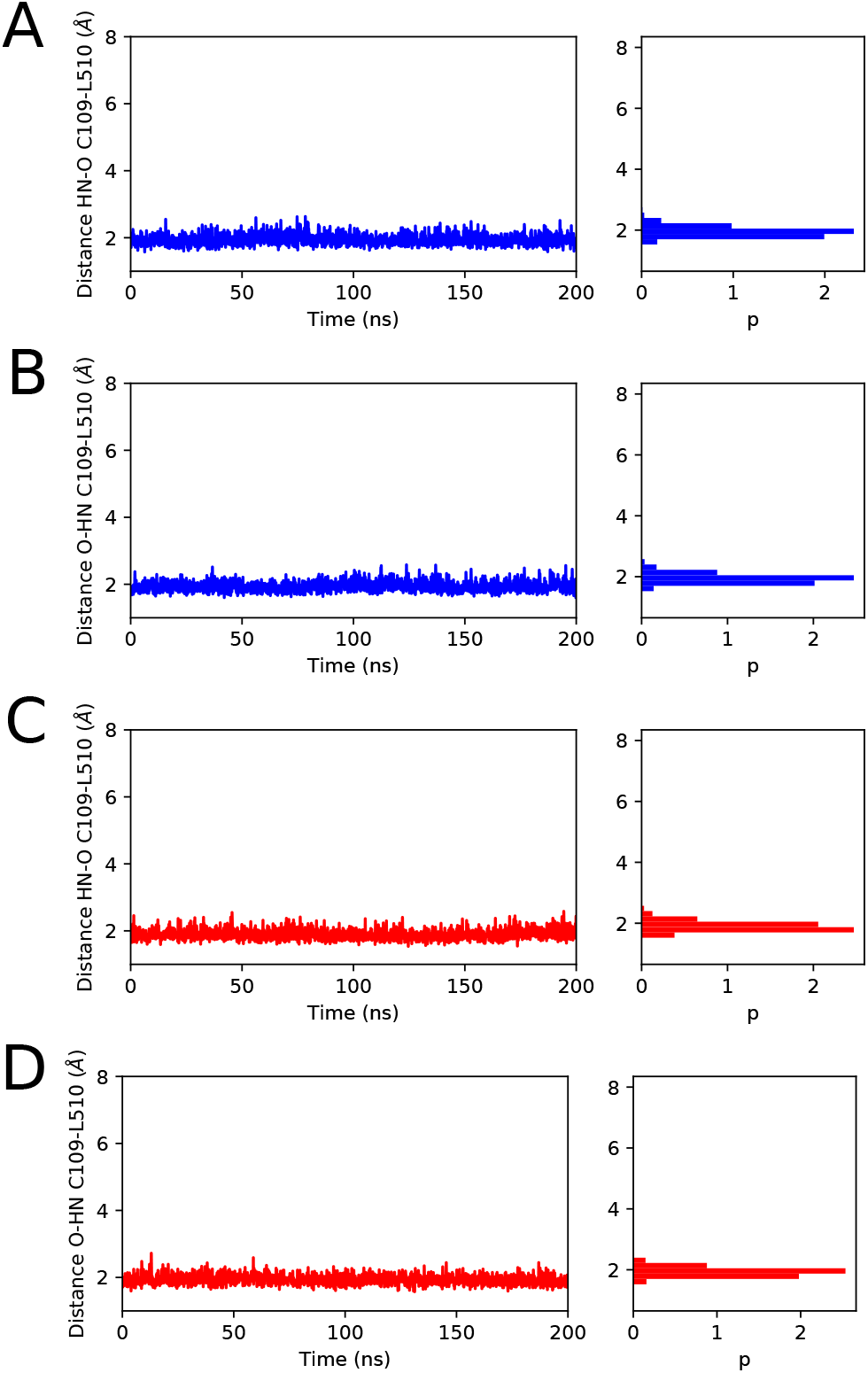
Tracking inter-domain hydrogen bonds in the simulations of cDsbD_red_-nDsbD_ox_ (A,B) and cDsbD_ox_-nDsbD_red_ (C,D).

The same is observed for C109 O L510 N and C109 O and L510 hydrogen bonds were maintained in 2 μs repeat simulations with the CHARMM36 force^37^ fieldS10. The O G107 to NH F531 hydrogen bond is more stable in the cDsbD_red_-nDsbD_ox_ than in the cDsbD_ox_-nDsbD_red_ complex.

**Figure S10:**
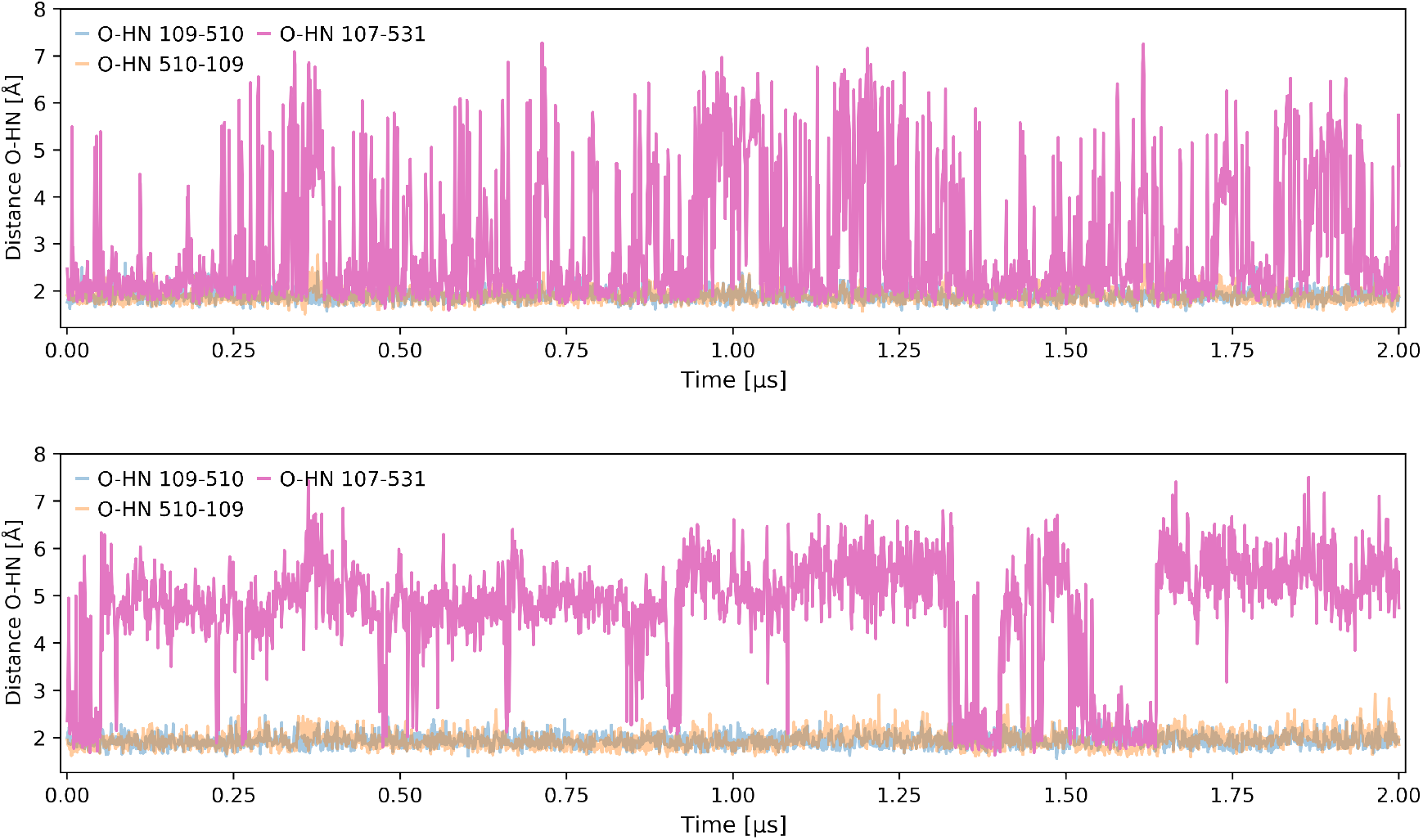
Repeat simulations of 2 μs with CHARMM36. (A) Stable cDsbD_red_-nDsbD_ox_ and (B) less stable cDsbD_ox_-nDsbD_red_ complexes.

#### Cap loop conformation in simulations of cDsbD-nDsbD complexes

The cap loop does not close in simulations of cDsbD_red_-nDsbD_ox_ and cDsbD_ox_-nDsbD_red_ (Fig. S11).

**Figure S11:**
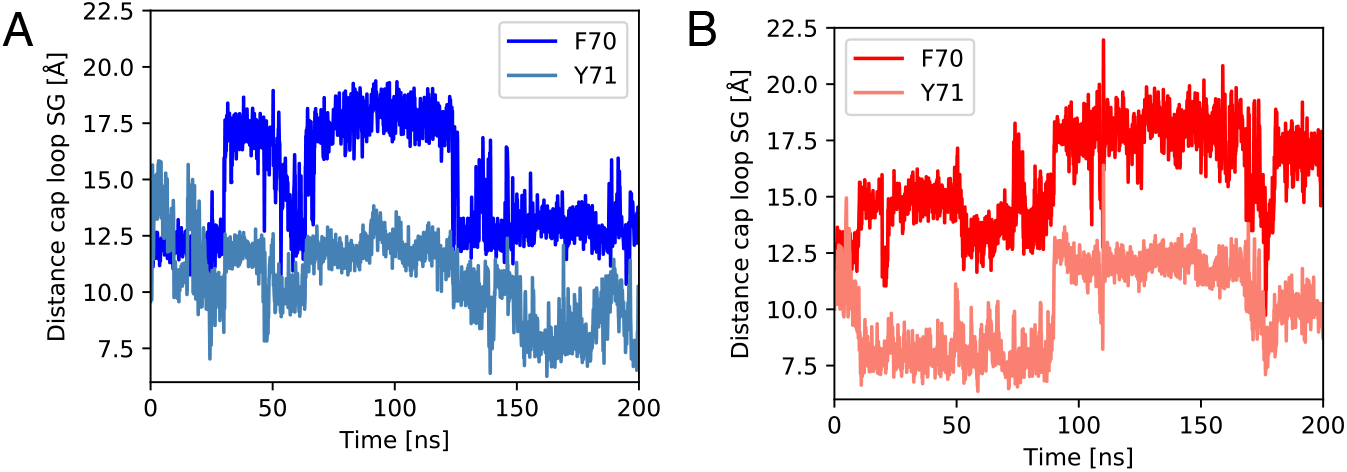
Following the cap-loop conformation in simulations of cDsbD_red_-nDsbD_ox_ (A) and cDsbD_ox_-nDsbD_red_ (B). Closed structures of the cap loop have F70-C109 distances 3-4 Åand Y71-C109 distances of 7 Å.^18^

### Additional AlphaFold modeling to investigate coupling between different domains and their conformations

To understand possible structural correlations between the different domains of full-length DsbD we re-run the model generation with either (1) cDsbD and linker or (2) nDsbD and linker removed (3) using a protocol designed to sample alternative conformations to investigate possible large-scale conformational changes in the transmembrane domain.

**Figure S12:**
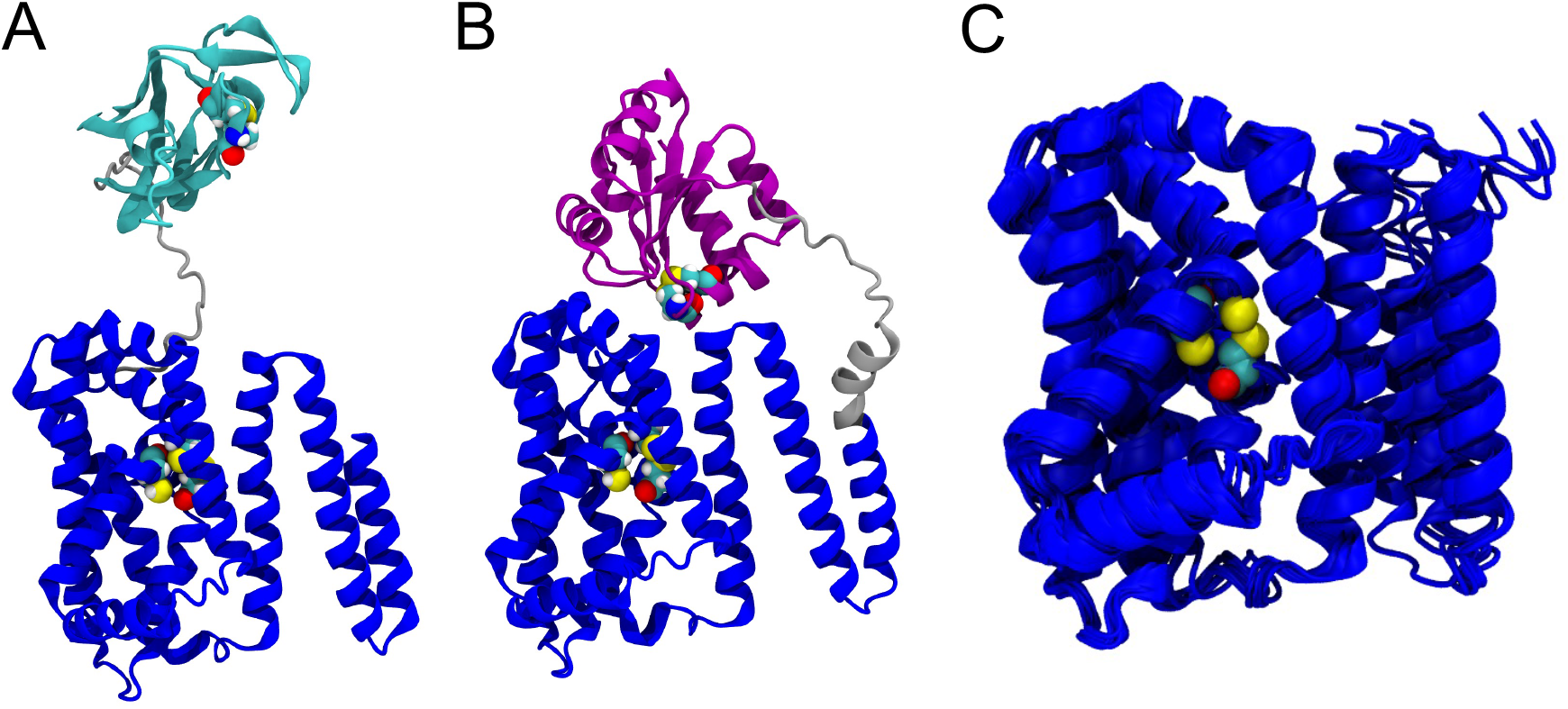
Understanding structural correlations between domains using AlphaFold modelling. (A) AlphaFold model of nDsbD and tmDsbD (residues 1-408) without cDsbD. (B) AlphaFold model of cDsbD and tmDsbD without nDsbD (residues 145-546). (C) Eight AlphaFold models of the transmembrane domain (residues 145-419) generated with the protocol of del Alamo *et al*.^41^ The structures are highly similar, with a maximum CA RMSD 145-408 to the average structure of the eight structures is 1.1 Å.

